# The Ca^2+^-gated Cl^-^ channel TMEM16A amplifies capillary pericyte contraction reducing cerebral blood flow after ischemia

**DOI:** 10.1101/2022.02.03.479031

**Authors:** Nils Korte, Zeki Ilkan, Claire Pearson, Thomas Pfeiffer, Prabhav Singhal, Jason Rock, Huma Sethi, Dipender Gill, David Attwell, Paolo Tammaro

## Abstract

Pericyte-mediated capillary constriction decreases cerebral blood flow in stroke after an occluded artery is unblocked. The determinants of pericyte tone are poorly understood. We show that a small rise in cytoplasmic Ca^2+^ concentration ([Ca^2+^]_i_) in pericytes activates chloride efflux through the Ca^2+^-gated anion channel TMEM16A, thus depolarizing the cell and opening voltage-gated calcium channels. This mechanism strongly amplifies the pericyte [Ca^2+^]_i_ rise and capillary constriction evoked by contractile agonists and ischemia. In a rodent stroke model, TMEM16A inhibition slows the ischemia-evoked pericyte [Ca^2+^]_i_ rise, capillary constriction and pericyte death, reduces neutrophil stalling and improves cerebrovascular reperfusion. Genetic analysis implicates altered TMEM16A expression in poor patient recovery from ischemic stroke. Thus, pericyte TMEM16A is a crucial regulator of cerebral capillary function, and a potential therapeutic target for stroke and possibly other disorders of impaired microvascular flow, such as Alzheimer’s disease and vascular dementia.

## INTRODUCTION

Cerebral blood flow (CBF) is regulated both at the arteriole and at the capillary level (1, 2), indeed capillaries are the site of highest vascular resistance within the brain (3, 4). Capillary resistance can be altered by changes in the tone of contractile pericytes with processes running circumferentially around the capillaries (5, 6). Electrical stimulation, optogenetically-induced depolarisation, contractile agonist application and optical ablation of single cortical pericytes have demonstrated the capacity of pericytes throughout the capillary bed to contract and control capillary diameter and local blood flow (7–13).

Alterations in pericyte contraction are crucial in the pathogenesis of ischemic stroke (6), Alzheimer’s disease (11), spreading depolarisation (e.g. during migraine with aura) (12) and neurological problems following cardiac arrest (13). After ischemic stroke, when blood flow to the occluded artery is restored, capillaries remain under-perfused (14, 15) even when thrombolysis is initiated shortly after stroke onset (16). This “no-reflow phenomenon” impairs patient recovery (17). The lack of reflow is largely due to pericytes constricting capillaries during and after ischemia (6, 18), possibly as a result of the cytoplasmic Ca^2+^ concentration ([Ca^2+^]_i_) rising in pericytes after a fall of ATP level ([ATP]i) inhibits ion pumping, although release of vasoconstrictors, such as endothelin-1 (ET-1) and thromboxane, in ischemic stroke may also contribute (19–21). Long-term capillary constriction will dramatically reduce local oxygen and glucose delivery, further aggravating the ischemic damage (22). For severe ischemia, pericyte-evoked capillary constriction is followed by pericytes dying in rigor (6, 18), thus prolonging the decrease of CBF. This pericyte loss (23), which is partly caused by the [Ca^2+^]_i_ rise that triggers contraction (6, 18), damages the blood-brain barrier (24–27). Neutrophil stalling in capillaries, which may be favored by pericyte-mediated narrowing of the capillary lumen, is also a proposed contributing factor to the no-reflow phenomenon after stroke (16, 28).

Preventing pericyte contraction could be of clinical benefit, but the determinants of pericyte tone are poorly understood. Pericyte contractility is controlled by [Ca^2+^]_i_ (29–31), which can be raised by depolarization activated Ca^2+^ (Cav) channels, or by Gq protein-coupled receptors (G_q_PCRs) triggering Ca^2+^ release from intracellular stores. However, Ca^2+^-activated Cl^-^ channels (specifically TMEM16A (32–34)) are found in smooth muscle and pericytes (35–38), and may be activated by any [Ca^2+^]_i_ rise that triggers contraction. Since smooth muscle cells (SMCs) and pericytes have a high intracellular Cl^-^ concentration (set by the plasma membrane Na^+^-K^+^-2Cl^−^ co-transporter NKCC1 and Cl^−^/HCO_3_^−^ exchanger AE2 (35–37)), when Cl^−^ channels open the resulting Cl^−^ efflux will cause a depolarization (for example, in kidney pericytes (37), the Nernst potential for Cl^-^ is around −30 mV). This depolarization is expected to activate Cav channels and amplify the increase in [Ca^2+^]_i_ and contraction that occur (39). In this paper, we test the hypotheses that (i) TMEM16A is a depolarising force in cerebral pericytes during agonist stimulation and ischemia and that (ii) inhibition of this channel opposes capillary constriction thus reducing tissue damage during ischemia.

From experiments in brain slices and in vivo, we demonstrate that TMEM16A is a crucial amplifier of cortical pericyte contraction evoked by [Ca^2+^]_i_ rises triggered by physiological modulators and ischemia. The importance of this is emphasised by a genetic analysis that implicates TMEM16A expression level as a determinant of recovery after stroke. Pharmacological inhibition of TMEM16A reduced the ischemia-evoked contraction and death of pericytes, improved post-ischemic CBF, decreased capillary neutrophil blocks at pericyte somata and reduced brain hypoxia and infarct size after ischemia, thus highlighting TMEM16A inhibition as a therapeutic strategy to improve reflow after stroke and other conditions of impaired microvascular blood flow.

## RESULTS

### TMEM16A generates Ca^2+^-activated Cl^-^ currents in cortical pericytes

TMEM16A mRNA is strongly expressed in cerebral pericytes and smooth muscle cells (38, 40), but the presence of TMEM16A protein in cerebral pericytes has not been previously assessed. We examined TMEM16A expression in pericytes of rat cortical slices, and in human cortical tissue removed surgically to access underlying tumours. Pericytes were labeled with antibodies to the proteoglycan NG2 (which is expressed by pericytes (41)) and TMEM16A, and the basement membrane around pericytes was visualised using isolectin B4 conjugated to an Alexa dye. Consistent with transcriptome studies (Supplementary Figure 1A), 90% of the TMEM16A expression was in pericytes (Figure 1A-B, Supplementary Figure 1B). Smooth muscle cells also expressed TMEM16A (Supplementary Figure 1C).

**Figure 1:**
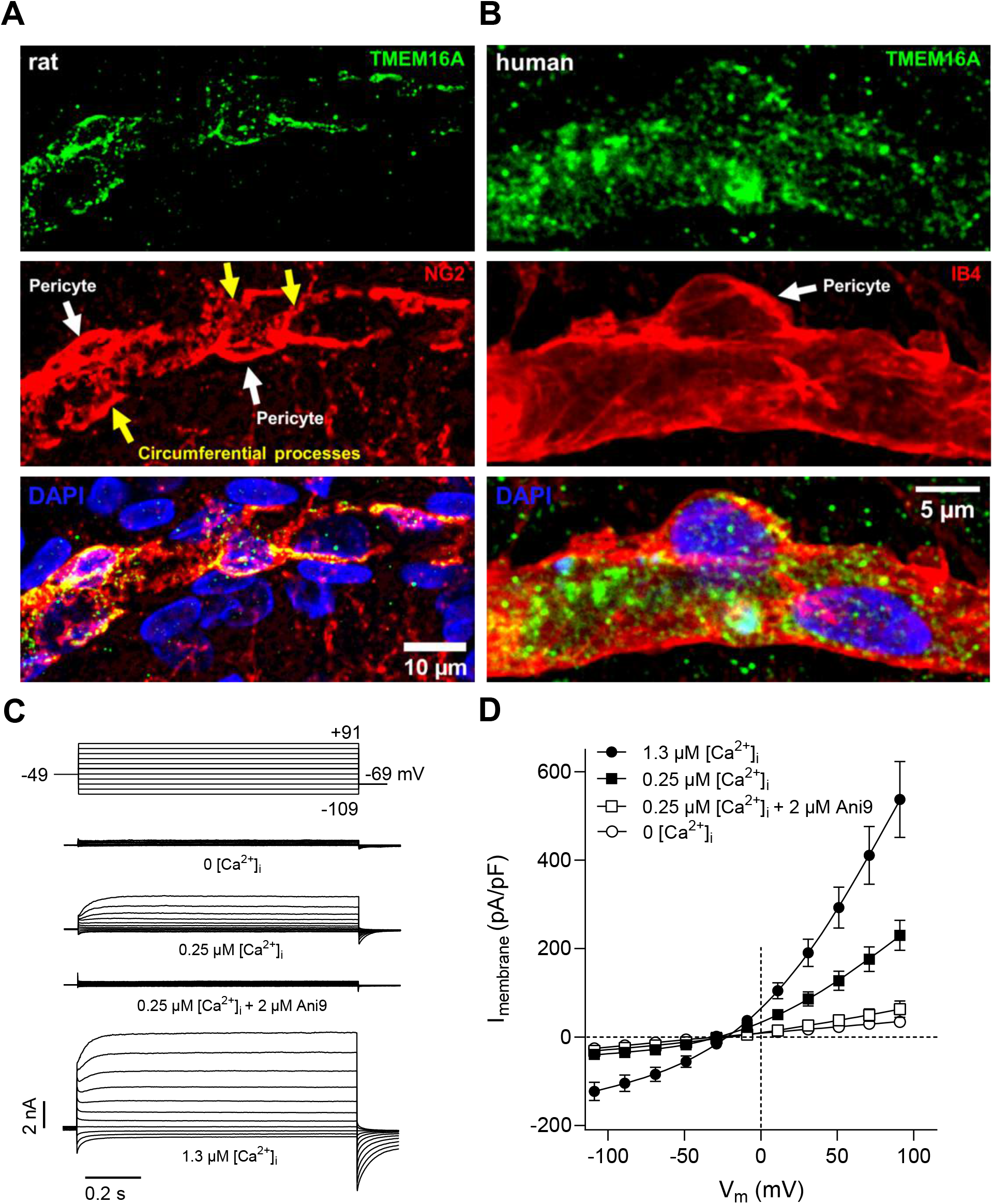
Cortical pericytes express functional TMEM16A channels. **(A)** TMEM16A expression in the soma and circumferential processes (yellow arrows) of NG2 labelled pericytes (white arrows) in a fixed cortical slice of a P21 rat. **(B)** TMEM16A expression in a pericyte labeled using isolectin B4 (IB4) in a fixed cortical slice of a 40-year-old human (representative of data from 5 subjects). **(C)** Representative family of whole-cell TMEM16A currents recorded from individual rat cortical pericytes using pipette solutions designed to isolate Cl^-^ currents, and with various free [Ca^2+^]_i_ (nominally 0, 0.25, or 1.3 μM), in the absence or presence of Ani9 (2 μM). The voltage protocol is illustrated in the upper panel, with the liquid junction potential corrected for. **(D)** Mean whole-cell TMEM16A current density versus voltage relationships in cortical pericytes (n=9-14) with various [Ca^2+^]_i_ and in the absence or presence of Ani9, as indicated.

To assess the importance of TMEM16A, pericytes were whole-cell clamped in rat cortical slices, with a Cs-based internal solution favouring recording of Cl^-^ currents (see Methods). With an intracellular solution containing 0.25 μM free [Ca^2+^]_i_, the steady-state current-voltage relationship was outwardly rectifying (Figure 1C-D), with a reversal potential (−27.2±4.2 mV, n=14) near the value of E_Cl_ (−25 mV). When [Ca^2+^]_i_ was raised to 1.3 μM, the membrane current either side of E_Cl_ was increased, and its rectification was reduced as observed for cloned TMEM16A channels (42, 43). With 0.25 μM [Ca^2+^]_i_, the TMEM16A inhibitor Ani9 (2-(4-Chloro-2-methylphenoxy)*N*’-(2-methoxybenzylidene)acetohydrazide) reduced the current (by 73% at 100 mV, p=0.02, Kruskal Wallis test with Dunn’s multiple comparisons test) to a level only slightly larger than when using Ca^2+^-free intracellular solution, for which TMEM16A channels should be closed (Figure 1D). Near the pericyte resting potential (~-40 mV), with 0.25 or 1.3 μM [Ca^2+^]_i_, Ca^2+^-activated Cl^-^ channels contributed a conductance of ~5 or 21 nS, respectively, to the cell. This is larger than the cell conductance with a normal K^+^-based internal solution and nominally zero [Ca^2+^]_i_ (~0.9 nS, see below), implying that TMEM16A has considerable scope to alter the cell’s membrane potential.

### Pericyte contraction evoked by G_q_PCR activation requires Ca^2+^ entry via Cav channels

Pericytes in acute rat cortical slices constricted capillaries in response to the contractile agonists ET-1 and U46619 (a thromboxane A2 analogue) which act on G_q_PCRs (9, 11). The circulating levels of ET-1 and thromboxane A2 are increased during stroke (19–21, 44). The capillary diameter at pericyte somata was reduced by ~70% (p<0.0001, paired two-tailed Student’s T-test) and ~20% (p<0.0001, paired two-tailed Student’s T-test) by ET-1 (10 nM) and U46619 (200 nM), respectively (Figure 2A-D).

**Figure 2:**
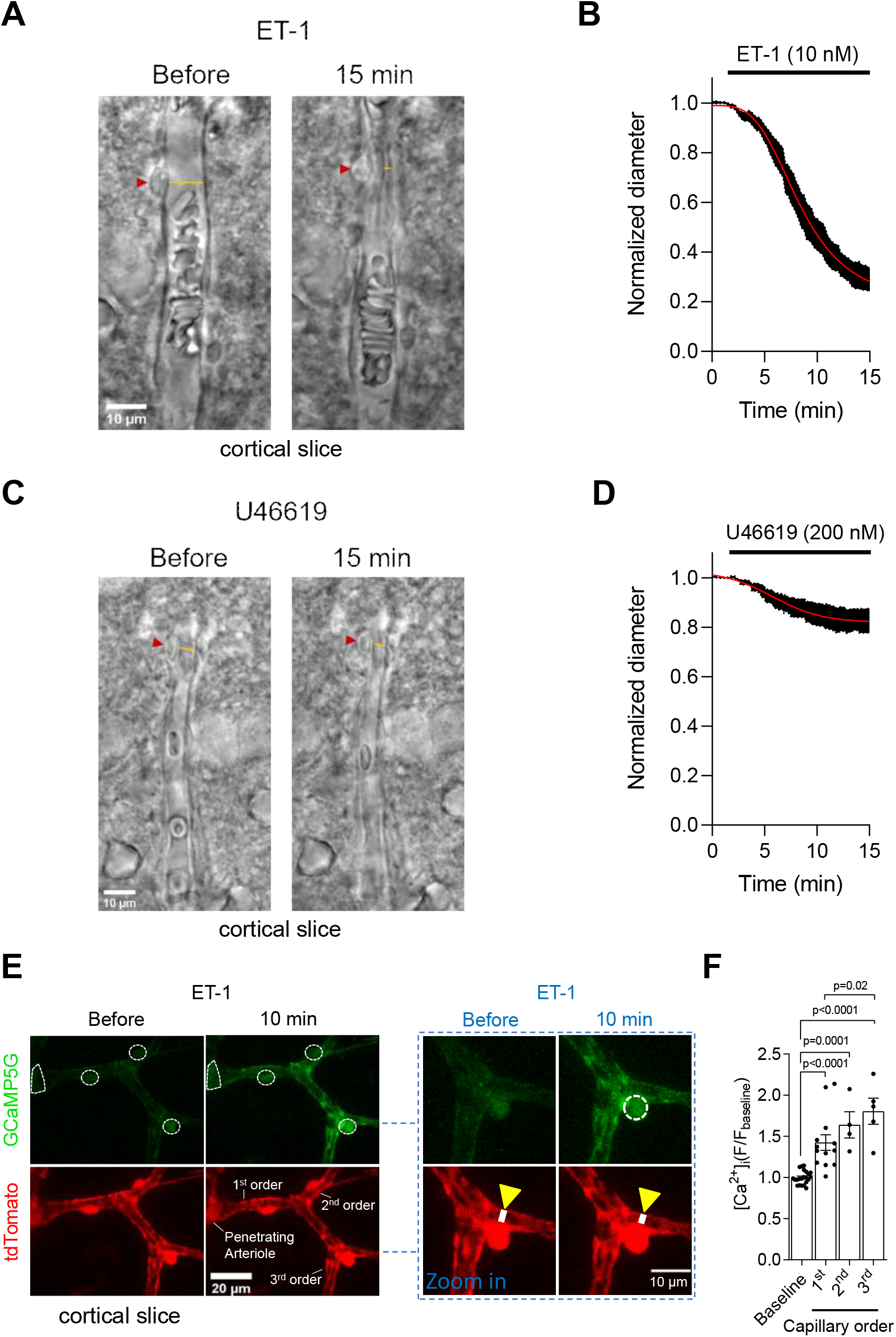
Vasoconstricting G_q_PCR agonists raise pericyte [Ca^2+^]_i_ and constrict capillaries at pericyte somata in acute cortical slices. **(A**) Representative bright-field images of a live rat cortical capillary pericyte before and after 15 min exposure to endothelin 1 (ET-1; 10 nM). Red arrowheads indicate the pericyte soma, and yellow lines indicate where the internal capillary diameter was measured. **(B)** Mean internal capillary diameter at pericyte somata during exposure to ET-1 (10 nM), normalized to the diameter measured in the absence of ET-1 (n=10). The ET-1 evoked capillary constriction was not dependent on the sex of rats (Supplementary Figure 3A). **(C)** Representative bright-field images of a live capillary pericyte as in (A). The thromboxane A2 analogue U46619 (200 nM) was applied as indicated. **(D)** Mean internal capillary diameter at pericyte somata during exposure to U46619 (200 nM), normalized to the diameter measured in the absence of U46619 (n=8). **(E)** Two-photon microscopy images (maximum intensity projections) of SMCs on a PA and pericytes on 1^st^-3^rd^ order capillary branches in acute cortical slices obtained from NG2-Cre^ERT2^-GCaMP5G mice. ET-1 raised the somatic [Ca^2+^]_i_ of SMCs and pericytes (encircled with white dashed lines). The pericyte [Ca^2+^]_i_ rise coincides with capillary constriction as indicated by the white line across the vessel lumen in the higher magnification image on the right. **(F)** ET-1 significantly raises [Ca^2+^]_i_ in 1^st^-3^rd^ order pericyte somata and evoked the greatest [Ca^2+^]_i_ rise in 3^rd^ order pericytes. The mean GCaMP5G fluorescence (F) in pericyte somata (points indicate individual pericytes from 5 mice) was normalized to the mean GCaMP5G fluorescence of the 17 min baseline (F_baseline_) with aCSF (One-way ANOVA with Tukey’s post hoc test).

The [Ca^2+^]_i_ rise evoked by ET-1 in pericytes was assessed using two-photon imaging of mice expressing tdTomato and the Ca^2+^ indicator GCaMP5G driven by the promoter for NG2 (NG2-Cre^ERT2^-GCaMP5G mice, see Methods). ET-1 increased [Ca^2+^]_i_ in pericytes on (at least) the 1^st^, 2^nd^ and 3^rd^ branch order capillaries from penetrating arterioles (PAs), where 1^st^ order refers to the first branch off the PA, 2^nd^ order refers to a branch off the 1^st^ order, etc. (Figure 2E-F). The [Ca^2+^]_i_ response to ET-1 increased significantly from the 1^st^ to the 3^rd^ branch order pericytes (Figure 2F). We show below that these differences in [Ca^2+^]_i_ rise are not due to a different ET-1 receptor response *per se*, but may reflect different numbers of TMEM16A and/or Cav channels in pericytes of different branch orders.

Ca^2+^ released from stores by G_q_PCR stimulation may activate actomyosin directly or may indirectly (via activation of TMEM16A) evoke depolarisation and thus activate Cav channels, causing an influx of extracellular Ca^2+^ and myofilament contraction. To distinguish between these possibilities, we applied ET-1 in the absence of extracellular Ca^2+^ ([Ca^2+^]_o_). In 0 [Ca^2+^]_o_, ET-1 neither significantly raised [Ca^2+^]_i_ (Figure 3A-C), nor evoked pericyte contraction (Figure 3D-E). Re-introducing [Ca^2+^]_o_ with ET-1 present rapidly raised [Ca^2+^]_i_ (Figure 3B-C) in the pericyte soma (p=6.2 x 10^-6^, (paired two-tailed Wilcoxon test with continuity correction) and processes (p<2.2 x 10^-16^ (paired two-tailed Wilcoxon test with continuity correction), presumably via extracellular Ca^2+^ entry (see below). This [Ca^2+^]_i_ rise evoked a significant capillary constriction at the soma of the pericytes (p=0.04, paired two-tailed Wilcoxon test with continuity correction) (Figure 3D-E), where most circumferential processes are located (10), but not at 10 μm (p=0.07, paired two-tailed Student’s t-test) or 20 μm (p=0.5, paired two-tailed Student’s t-test) along the capillary from the pericyte soma (Figure 3E), where the processes run more longitudinally (11). Capillary diameter was larger at the soma than away from the soma (Figure 3E) in the absence of ET-1 (p=0.002, unpaired two-tailed Student’s t-test) and also after 15 min in 0 [Ca^2+^]_o_ and ET-1 (p=0.003, unpaired two-tailed Student’s t-test), consistent with measurements in vivo (6), suggesting that the pericyte soma may release factors that induce growth of the endothelial tube.

**Figure 3:**
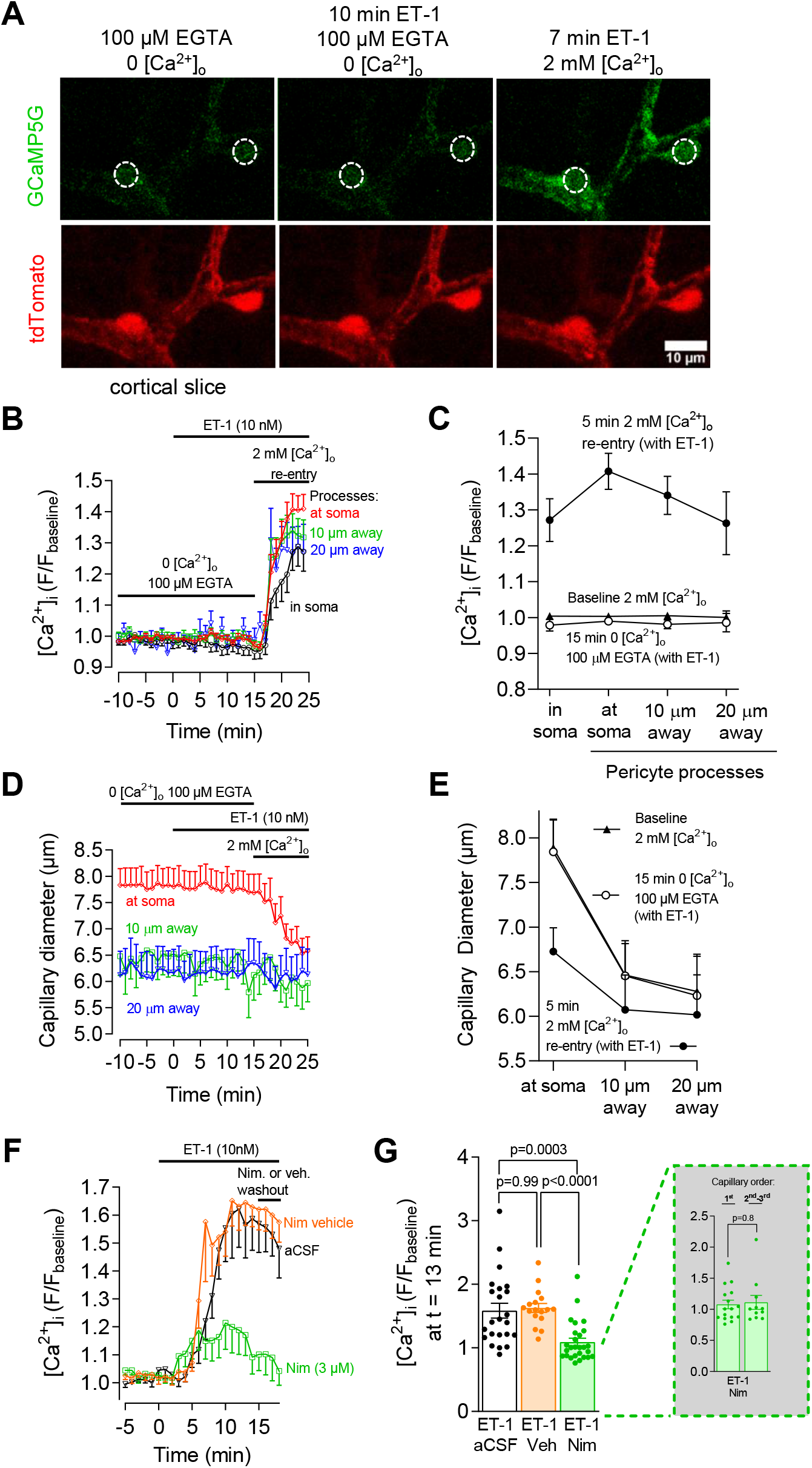
Pericyte contraction evoked by G_q_PCR activation requires Ca^2+^ entry via Cav channels. **(A)** Two-photon microscopy images (maximum intensity projections) of pericytes on 1^st^-3^rd^ order capillary branches from the PA in an acute cortical slice of a NG2-Cre^ERT2^-GCaMP5G mouse. Dashed white circles are pericyte somata. **(B)** Removing extracelluar Ca^2+^ abolished the ET-1 evoked [Ca^2+^]_i_ rise. Time course of GCaMP5G fluorescence (F) in pericyte somata (red trace; n=32) and processes (other traces; n=79) normalized to mean baseline fluorescence with 2 mM [Ca^2+^]_o_ (the last 10 min of 15 min in 0 [Ca^2+^]_o_ are shown). [Ca^2+^]_i_ changes in processes were quantified <5 μm from pericyte somata centers (“at soma”), and at 10 μm and 20 μm along the vessel from the soma center. **(C)** 15 min of 0 [Ca^2+^]_o_ did not affect pericyte [Ca^2+^]_i_ (compare bottom two plots). Re-introducing Ca^2+^ in the continuous presence of ET-1 raised pericyte [Ca^2+^]_i_ in the processes and somata (see also (B)). **(D)** Capillary constriction at pericyte somata (n=40) coincides with the [Ca^2+^]_i_ rise upon 2 mM [Ca^2+^]_o_ re-perfusion in (B). There was no significant change in capillary diameter away from pericyte somata at 10 (n=18) or 20 μm (n=11). **(E)** Capillary diameter is larger at baseline, and constricts in response to 2 mM [Ca^2+^]_o_ re-perfusion, at pericyte somata. In (D) and (E) diameter is from tdTomato channel. **(F)** Time-course of ET-1 evoked [Ca^2+^]_i_ change in pericyte somata, normalized to aCSF baseline. Nimodipine (3 μM) or vehicle were applied 15 min prior to ET-1 application. **(G)** Nimodipine greatly attenuated the ET-1 evoked pericyte [Ca^2+^]_i_ rise (note that zero [Ca^2+^]_i_ is at 1 on y-axis) (Kruskal Wallis test with Dunn’s post hoc test). *Inset* In nimodipine, the ET-1 evoked [Ca^2+^]_i_ rise was similar in pericytes on 1^st^ order versus 2^nd^ to 3^rd^ order branches (unpaired two-tailed Student’s T-test).

To test whether the absence of an ET-1 evoked pericyte [Ca^2+^]_i_ rise and contraction in 0 [Ca^2+^]_o_ was the result of preventing Ca^2+^ entry via Cav channels, or alternatively a result of depleting internal stores of Ca^2+^, we examined the effect of the L-type Cav blocker nimodipine on the ET-1 response in normal [Ca^2+^]_o_ solution. Nimodipine (3 μM) inhibited the [Ca^2+^]_i_ rise by 86% (Figure 3F-G), implying that most Ca^2+^ enters via Cav channels (ET-1 can also activate, for example, TRPM4 and TRPC3, 5, 6 and 7 channels in other cell types, however the 86% suppression of the steady state [Ca^2+^]_i_ rise by nimodipine in Figure 3G implies that VGCCs contribute the great majority of the Ca^2+^ influx.). The [Ca^2+^]_i_ rise seen in nimodipine was similar in 1^st^ and 2^nd^-3^rd^ order pericytes (Figure 3G), suggesting that heterogeneity in the [Ca^2+^]_i_ rise between 1^st^-3^rd^ order pericytes (Figure 2F) may involve differences in the number of L-type Cav channels activated downstream of the small [Ca^2+^]_i_ rise caused by ET-1 receptors alone.

Together these data demonstrate that most of the ET-1 triggered contraction requires Ca^2+^ entry via Cav channels. This raises the question of how Ca^2+^ released from internal stores by ET-1 leads to Cav activation.

### TMEM16A amplifies the pericyte [Ca^2+^]_i_ rise and contraction evoked by vasoconstrictors

Two structurally unrelated TMEM16A inhibitors, MONNA (45) and Ani9 (46), did not affect the diameter of cortical capillaries when they were not exposed to exogenous constrictors (Figure 4A), consistent with the idea that TMEM16A channels have no or low activity in the absence of substantial G_q_PCR stimulation. Lack of basal TMEM16A activity may reflect a decrease in vascular tone in brain slices, possibly as a result of a lack of blood flow and the shear stress it generates, or lack of noradrenaline release from the axons of locus coeruleus neurons (which become severed in the brain slicing procedure). However, MONNA and Ani9 greatly reduced the capillary constriction evoked by ET-1 (10 nM) (Figure 4B) and U46619 (200 nM) (Figure 4C). Furthermore, Ani9 strongly attenuated the ET-1 evoked [Ca^2+^]_i_ rise in all tested capillary branch orders (Figure 4D). The 94% reduction of the ET-1 evoked [Ca^2+^]_i_ rise produced by Ani9 in Figure 4D, where fluorescence change is measured from the *F/F_baseline_* value of 1, is not inconsistent with the ~75% reduction of TMEM16A current in Figure 1D: there is no reason to expect a linear relationship between the suppression of the [Ca^2+^]_i_ rise and the suppression of the TMEM16A current, because of the non-linear relationships that exist between [Ca^2+^]_i_ and TMEM16A activation and between TMEM16A-evoked depolarization and voltage-gated Ca^2+^ current activation. These results are consistent with a TMEM16A-mediated depolarization amplifying the ET-1 evoked rise of [Ca^2+^]_i_ by activating Cav channels.

**Figure 4:**
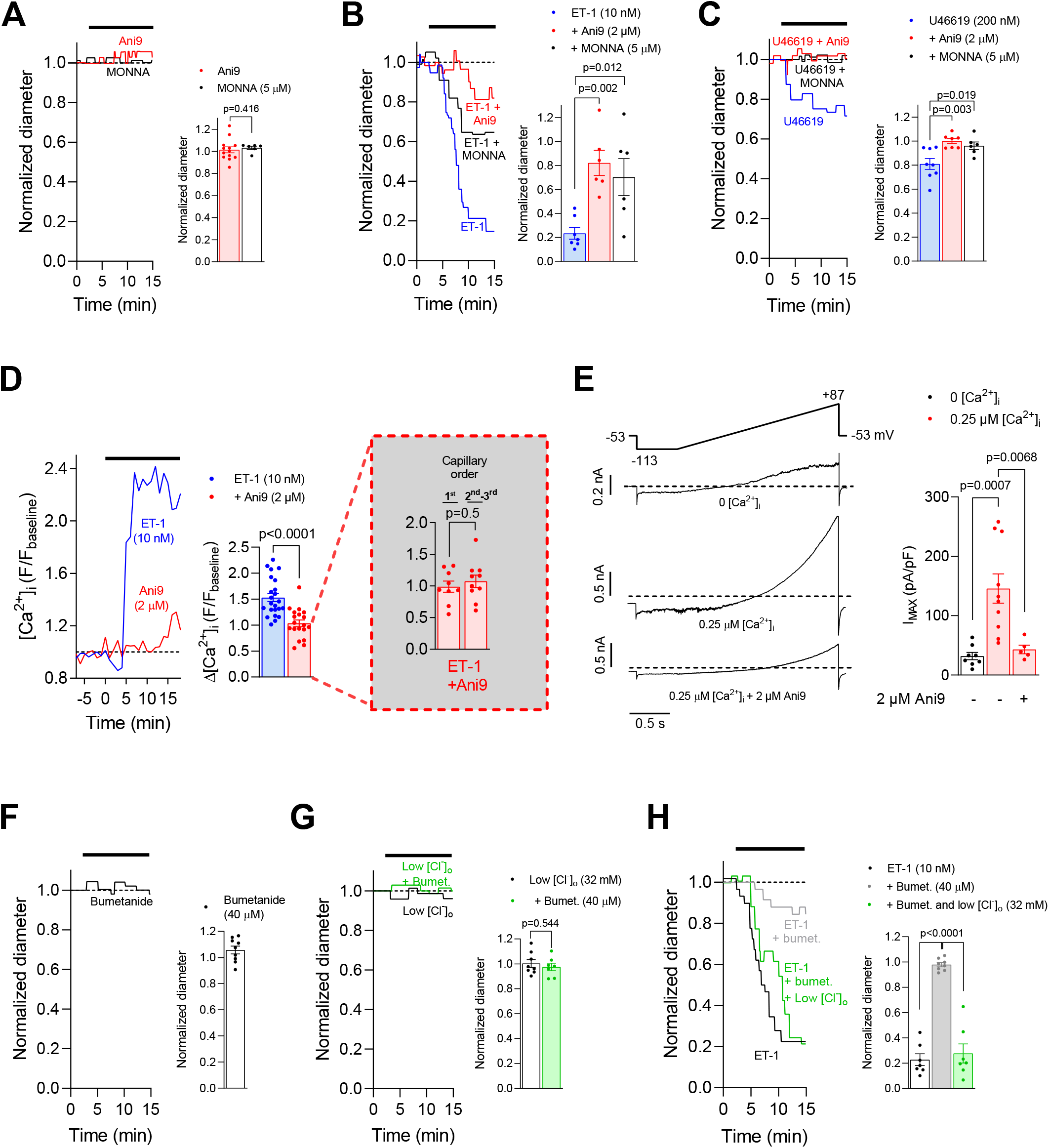
Effects of Cl^-^ on TMEM16A-mediated control of pericyte [Ca^2+^]_i_ and tone. **(A)** Left: capillary diameter at pericyte somata during Ani9 (2 μM) or MONNA (5 μM) superfusion in acute rat cortical slices, normalized to baseline diameter. Right: mean normalized capillary diameter after exposure to Ani9 or MONNA **(B-C)** Ani9 or MONNA reduced capillary constriction evoked by ET-1 (10 nM, B) or U46619 (200 nM, C). **(D)** Ani9 reduced the ET-1 evoked [Ca^2+^]_i_ rise in pericyte somata in cortical slices of NG2-Cre^ERT2^-GCaMP5G mice. GCaMP5G fluorescence (F) was normalized to mean baseline fluorescence in aCSF. *Inset* In Ani9, the ET-1 evoked [Ca^2+^]_i_ rise was similar in pericytes on 1^st^ order versus 2^nd^ to 3^rd^ order branches from the PA. **(E)** Whole-cell currents of rat cortical pericytes in acute rat cortical slices (left panel). Voltage protocol is shown with the liquid junction potential corrected for. Free [Ca^2+^]_i_ was 0 or 0.25 μM. Right panel shows the mean whole-cell current density (I_MAX_) at the end of the ramp (i.e. +87 mV) in 0 or 0.25 μM [Ca^2+^]_i_. **(F)** Normalized capillary diameter at the pericyte soma during bumetanide (40 μM) superfusion in an acute rat cortical slice (left panel). Right panel shows mean normalized capillary diameter at pericyte somata after exposure to bumetanide (40 μM). **(G)** Effects of low [Cl^-^]_o_, and bumetanide on rat cortical capillary diameter. **(H)** Changes in capillary diameter in response to ET-1 after 15 min pre-incubation with bumetanide, with or without a lower [Cl^-^]_o_. In G and H, left panels show representative capillary responses, and right panels show normalized capillary diameter. Number of animals are detailed in Suppl. Table 2. Statistical comparisons: A: Mann Whitney, D and G or unpaired two-tailed Student’s T-test; in B, C, E and H: one-way ANOVA with Bonferroni’s post hoc multiple comparisons test.

The Ani9-mediated reduction of the ET-1 evoked [Ca^2+^]_i_ rise and pericyte contraction is unlikely to result from Ani9 indirectly affecting pericyte tone by acting on nearby endothelial cells, astrocytes or neurons, since TMEM16A is nearly exclusively expressed in cortical pericytes and smooth muscle cells (Figure 1A-B, Supplementary Figure 1A-C) (47, 48). Furthermore, in whole-cell clamped pyramidal neurons, the action potential response to current injection, the resting potential, and the neuronal input resistance were not affected by Ani9 (Supplementary Figure 2).

We confirmed that TMEM16A is present at sufficient density to significantly change the cell’s membrane potential, and thus activate Cav channels, by whole-cell clamping pericytes with physiological internal solutions containing 0 or 0.25 μM free [Ca^2+^]_i_ (see Methods). With 0 free [Ca^2+^]_i_, the resting potential and input resistance were −42.0±4.5 mV and 1.1±0.5 GΩ respectively in 8 cells. With 0.25 μM [Ca^2+^]_i_, an inward current of −12.8±2.3 (n=10) pA/pF (mean cell capacitance: 10.6±1.0 pF (n=10)) was present at −40 mV (increased from a value of +0.3±1.2 pA/pF (n=8) in 0 [Ca^2+^]_i_ (mean cell capacitance: 10.3±1.0 pF (n=8)). This current was reduced to −5.3±0.7 (n=5) pA/pF in the presence of 2 μM Ani9 (mean cell capacitance: 11.4±1.6 pF (n=5)), implying a TMEM16A-mediated Cl^-^ conductance of ~5 nS. This Ca^2+^-activated current depolarized the resting potential to −18.3±4.1 mV (n=10) (Figure 4E).

### Pericyte tone is strongly influenced by the transmembrane Cl^-^ gradient

If TMEM16A confers a fundamental depolarizing mechanism recruited during G_q_PCR activation, altering the transmembrane Cl^-^ gradient should alter the amplification of constriction produced by TMEM16A. This gradient was altered in cortical slices using three different strategies: (i) exposing slices to bumetanide (40 μM) to inhibit the Cl^-^ importer NKCC1, and thus reduce the intracellular Cl^-^ concentration ([Cl^-^]_i_) and the depolarizing influence of TMEM16A; (ii) reducing the extracellular Cl^-^ concentration ([Cl^-^]_o_) to increase TMEM16A-mediated depolarization, or (iii) a combination of these treatments. In slices not exposed to a G_q_PCR agonist, alterations in the Cl^-^ gradient did not affect capillary diameter (Figure 4F-G), consistent with TMEM16A channels being closed under these conditions. In contrast, when pericytes were exposed to ET-1 following pre-treatment with bumetanide (to reduce the depolarizing [Cl^-^]_i_ gradient), the resulting capillary constriction was attenuated approximately 4-fold (Figure 4H). Lowering [Cl^-^]_o_ to re-instate a depolarising Cl^-^ gradient in the presence of bumetanide restored the ET-1 evoked capillary pericyte contraction (Figure 4H) to a magnitude similar to that seen with ET-1 application alone (Figure 4B). Thus, capillary pericyte contraction is strongly dependent on the transmembrane Cl^-^ gradient.

### TMEM16A KO and raising [K^+^]_o_ confirm that TMEM16A amplifies constrictions

Using wild-type mice or floxed TMEM16A (*ANO1*) knock-out (KO) mice crossed with NG2-Cre^ERT2^ mice (to knock-out TMEM16A in NG2-expressing pericytes), we assessed the effect of ET-1 on the pericyte-mediated capillary constriction. This showed that KO of TMEM16A reduced the ET-1 evoked constriction of capillaries by pericytes (Figure 5A-B) as was also seen when using pharmacological inhibition of TMEM16A. The smaller effect of the KO in Figure 5A, compared to pharmacological inhibition in Figure 4B, likely reflects the incomplete KO achieved with Cre (the reported percentage of cells, probably oligodendrocyte precursor cells, that undergo recombination was ~60% at 2 weeks after tamoxifen administration for the Cre line we used (49) although in retinal pericytes using a different NG2-Cre line (50) it was only ~30%).

**Figure 5:**
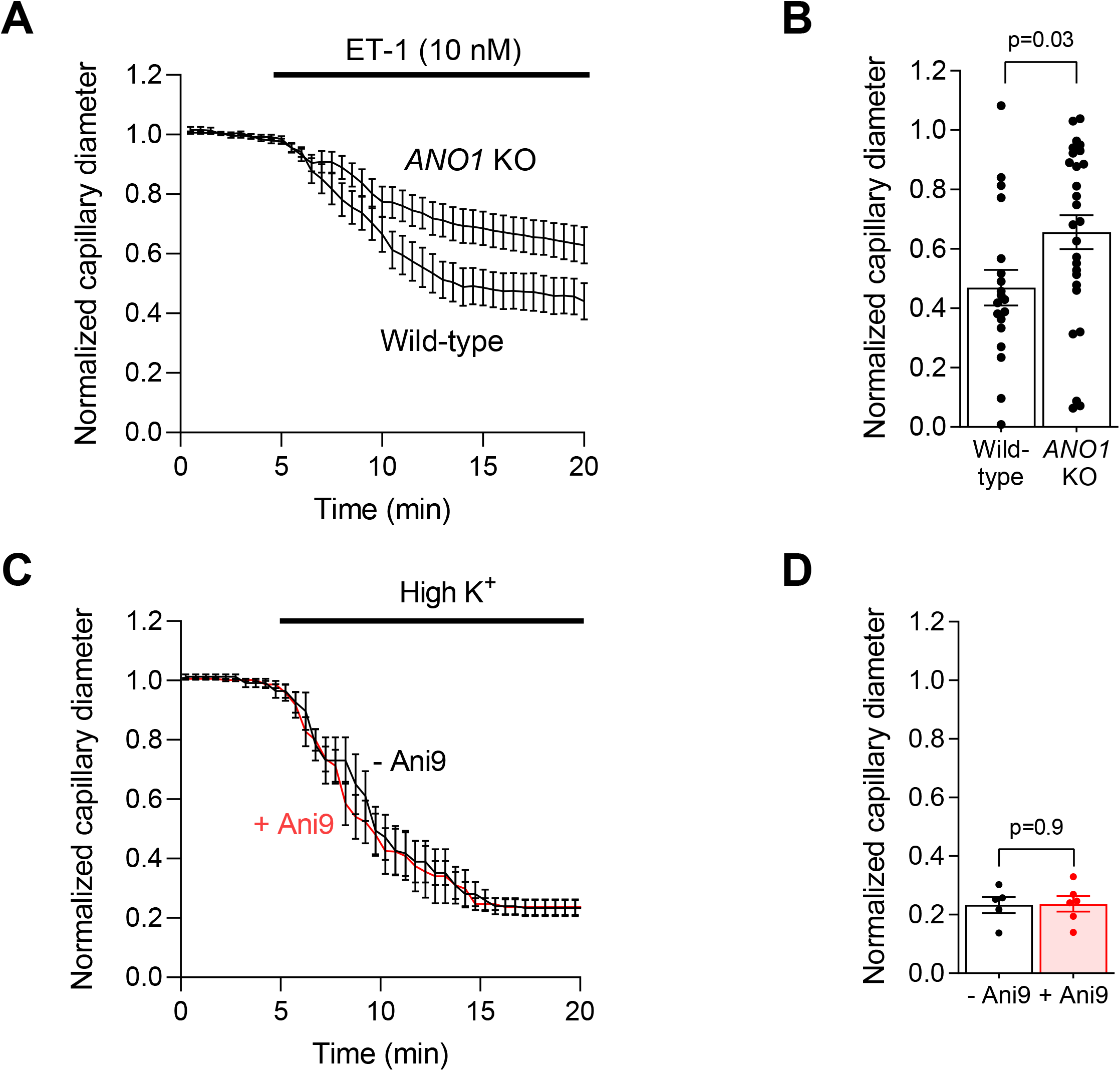
TMEM16A knock-out (KO) in pericytes reduces endothelin-1 (ET-1) evoked pericyte contraction and depolarizing the membrane potential induces pericyte contraction independent of TMEM16A block. **(A)** Mean internal capillary diameter at pericyte somata during exposure to ET-1 (10 nM), normalized to the diameter measured in the absence of ET-1, in acute cortical slices of wildtype or TMEM16A (*ANO1*) KO mice. **(B)** TMEM16A KO reduced the average ET-1 evoked pericyte contraction during the last 5 minutes of the experiment (Ano KO, n=27; Wild-type, n=19). **(C)** Mean internal capillary diameter at pericyte somata during exposure to 92.5 mM extracellular potassium ([K^+^]_o_), normalized to the diameter measured in the presence of 2.5 mM [K^+^]_o_ in acute rat cortical slices. **(D)** Raising [K^+^]_o_ evokes pericyte contraction, and this is not reduced by Ani9 (2 μM). Points indicate individual pericytes from 5 rats per condition. Statistical comparisons in B and D: Unpaired two-tailed Student’s T-test.

Depolarising the membrane potential in rat brain slices by raising the extracellular potassium concentration ([K^+^]_o_) to 92.5 mM, which is expected to activate voltage-gated Ca^2+^ channels without the need for TMEM16A activation, elicited a capillary constriction (Figure 5B-C) similar to that induced by application of ET-1 (Figure 2B). Consistent with this approach bypassing the intermediary step of TMEM16A activation, the constriction evoked by high [K^+^]_o_ was unaffected by Ani9 (Figure 5D).

### TMEM16A inhibition attenuates capillary constriction and pericyte death in ischemia

In ischemia, low [ATP]i will slow Ca^2+^ pumping out of pericytes, raising [Ca^2+^]_i_. This may activate TMEM16A and promote Cav activation. We used Ani9 to test whether blocking TMEM16A could reduce the pericyte contraction and death that ischemia evoke (6).

Capillary diameters and pericyte [Ca^2+^]_i_ were measured in acute cortical slices during oxygen and glucose deprivation (OGD), in the absence or presence of Ani9 (2 μM). Ani9 strongly delayed pericyte-mediated capillary constriction (Figure 6A-B), and reduced the OGD-evoked [Ca^2+^]_i_ rise by ~70% (Figure 6C-D). Since ischemia-evoked pericyte death is partly driven by Ca^2+^ influx (5), we tested whether TMEM16A inhibition affected OGD-induced pericyte death in rat cortical slices incubated for 1 hour in aCSF solution (control) or in OGD solution in the absence or presence of Ani9 (2 μM). Propidium iodide uptake was used as a marker of necrotic death of pericytes. Ani9 reduced the OGD-induced pericyte death from 50.1% to 28.7%, compared with 7.6% in normal aCSF (Figure 6E-F).

**Figure 6:**
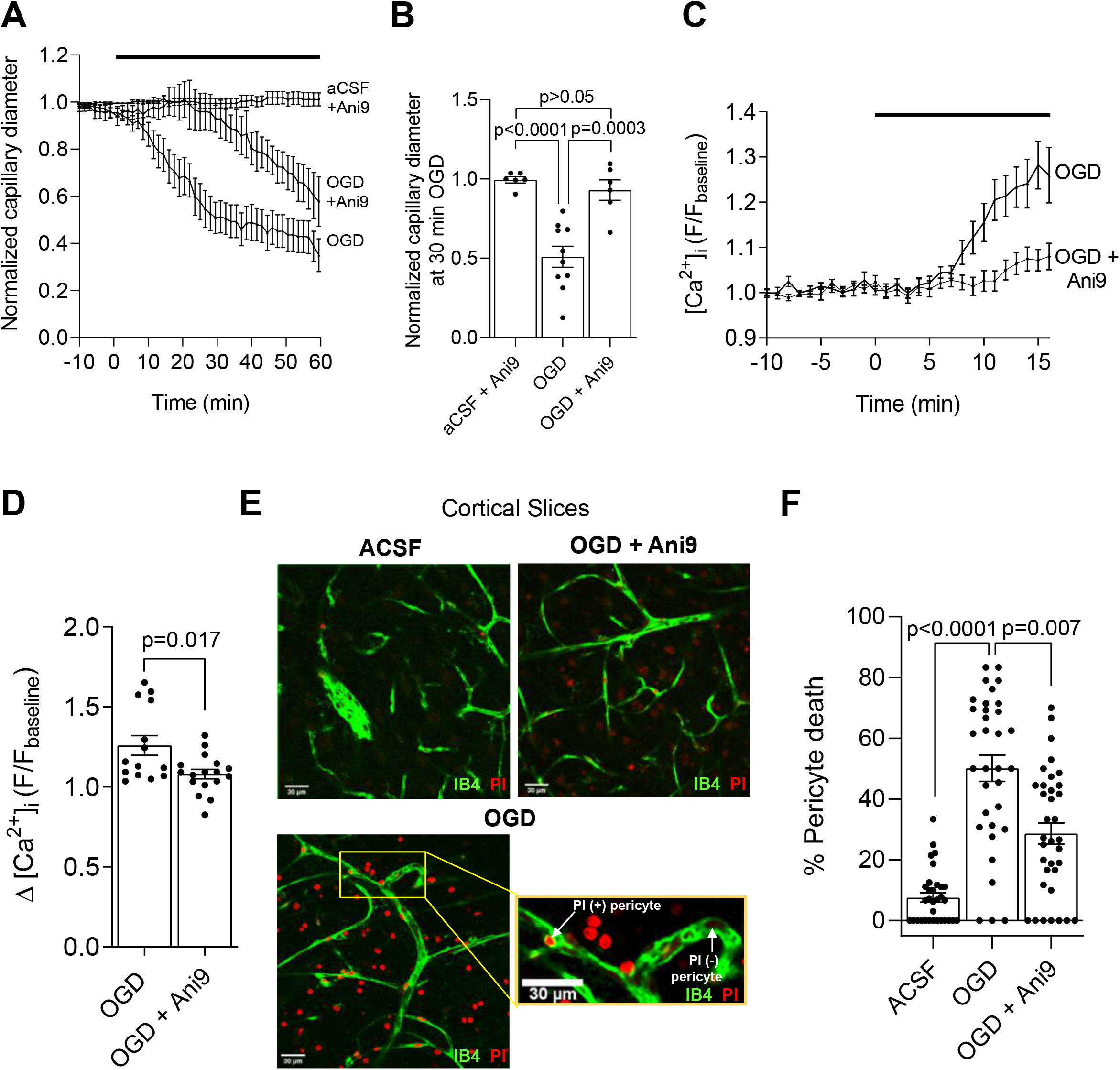
Blocking TMEM16A slows the ischemia-evoked [Ca^2+^]_i_ rise in pericytes, delays capillary constriction and reduces pericyte death in acute cortical slices. **(A)** Mean normalized capillary diameter at pericyte somata during perfusion with aCSF (control, n=6) or oxygen and glucose deprived (OGD) solution in the presence (n=6) or absence (n=8) of Ani9 (2 μM) in acute rat cortical slices. **(B)** Ani9 reduces the ischemia-evoked pericyte-mediated capillary constriction at 30 min of OGD. Points indicate individual pericytes from 5-6 rats per condition. The OGD evoked capillary constriction was not dependent on the sex of rats (Supplementary Figure 3B) (One-way ANOVA with Bonferroni’s post hoc multiple comparisons test). **(C)** Time course of the mean change in normalized GCaMP5G fluorescence (F) in pericyte somata during OGD with or without Ani9 (2 μM) in NG2-Cre^ERT2^-GCaMP5G mice. **(D)** Ani9 reduces the [Ca^2+^]_i_ rise (measured from the y-axis value of 1) at 16 min of perfusion with OGD. Points indicate individual pericytes from 3 mice per condition (Unpaired two-tailed Student’s T-test with Welch’s correction). **(E)** Confocal images of rat cortical capillaries labelled with isolectin B4 (IB4) to visualize pericytes labeled by the necrosis marker propidium iodide (PI) after 1 hr exposure to aCSF or OGD in the presence or absence of Ani9 (2 μM). In the lower panel, the inset illustrates examples of necrotic (PI +) and healthy (PI -) pericytes. **(F)** Ani9 reduces the OGD-evoked pericyte death. The percentage of dead pericytes was quantified by dividing the number of PI-labelled pericytes by the total number of pericytes in images as shown in (E) (Kruskal Wallis test with Dunn’s post hoc test). Number of animals are detailed in Suppl. Table 2.

### Genetic evidence for a role of TMEM16A in stroke

The experiments above implicate TMEM16A in the control of capillary diameter in response to G_q_PCR stimulation and ischemia, and pericyte death in ischemia. These observations suggest an involvement of TMEM16A in the capillary constriction that follows ischemic stroke. To investigate this in human subjects, we identified a genetic proxy for TMEM16A activity (Table 1) and applied this in a Mendelian randomization analysis (51) to explore its association with (i) the risk of developing ischemic stroke (using the MEGASTROKE genetic association study (52)) and (ii) recovery after ischemic stroke (using the GISCOME study (53)). The genetic proxy for TMEM16A activity was a single nucleotide polymorphism (rs755016, in an intron of the *Ano1* gene that encodes TMEM16A) that associated with (i) *TMEM16A* gene expression (based on data from the GTEx database (54)), and (ii) raised diastolic blood pressure (based on a study of 757,601 participants: Supplementary Table 1 of Ref. (55)). Diastolic blood pressure was used to select the genetic proxy for TMEM16A activity because TMEM16A is known to increase systemic vascular resistance and blood pressure (56, 57). Phenome-wide association study of this proxy predominantly identified associations with psychiatric, neurological and circulatory traits (Supplementary Table 1), and the other traits identified may also have a circulatory component.

**Table 1.** Single-nucleotide polymorphism (SNP) used as a proxy for TMEM16A activity in Mendelian randomization analyses. The strongest association with *TMEM16A* gene expression was observed in thyroid tissue (*P*=4.8×10^-14^). A strong association was also observed in the meta-analysis considering all available tissues (*P*=2.73×10^-7^), showing that this proxy was not only specific to TMEM16A in the thyroid. The association with diastolic blood pressure (P=9.81×10^-6^) suggests that this variant also serves as a proxy for TMEM16A activity, and not just gene expression.

| SNP ID | Chr | Pos<br>(hg19) | EA | OA | EAF | Increasing<br>*ANO1*<br>expression<br>in thyroid | Increasing<br>*ANO1*<br>expression<br>across all<br>tissues | Diastolic<br>blood<br>pressure | Stroke risk<br>(log OR) |  |  | Stroke recovery<br>(log OR; positive estimates<br>indicate worse outcome,<br>mRS 3-6 rather than 0-2) |  |  |
| --- | --- | --- | --- | --- | --- | --- | --- | --- | --- | --- | --- | --- | --- | --- |
|  |  |  |  |  |  | *P* | *P* | *P* | Beta | SE | *P* | Beta | SE | *P* |
| rs755016 | 11 | 69952562 | G | A | 0.59 | 4.80E-14 | 2.73E-07 | 9.81E-06 | 0.01 | 0.01 | 0.25 | 0.13 | 0.05 | 0.01 |

*Chr: chromosome; Pos: position; hg19: human genome reference panel used to annotate variants; EA: effect allele; OA: other allele; EAF: effect allele frequency; OR: odds ratio; Beta: association estimate from GWAS; SE: standard error*

Our analysis showed that, while genetically proxied TMEM16A activity was not associated with increased risk of ischemic stroke (odds ratio [OR] per allele associated with increased *TMEM16A* expression 1.01, 95% confidence interval [CI] 0.99-1.03, p=0.25), increased genetically proxied *TMEM16A* expression associated with worse functional outcome (score 3-6 of the modified Rankin Scale) at 60-190 days following ischemic stroke (OR 1.13, 95% CI 1.03-1.25, p=0.01). The association of genetically proxied TMEM16A activity with outcome after ischemic stroke, but not with risk of ischemic stroke, makes effects on systemic blood pressure an unlikely mediator, as this affects ischemic stroke risk more than recovery (58). Searching the PhenoScanner database for disease outcomes or traits related to the genetic variant proxying TMEM16A activity (59) revealed no genome-wide significant associations to suggest pleiotropic effects that could bias the Mendelian randomization analysis. Thus, this analysis suggests that increased *TMEM16A* expression is associated with a worse outcome after human ischemic stroke. We therefore examined the influence of TMEM16A inhibition in an in vivo rodent stroke model.

### TMEM16A block decreases pericyte contraction in vivo and improves CBF after stroke

Bilateral common carotid artery occlusion (CCAO) was performed in mice to mimic the ischemia and reperfusion injury that occurs during severe ischemic stroke (60). Using simultaneous laser Doppler flowmetry and two-photon imaging in vivo, we tested the effect of TMEM16A inhibition on pericyte [Ca^2+^]_i_, capillary diameter and CBF after ~7.5 min of CCAO in the cortex (in vivo) of NG2-dsRed mice or NG2-Cre^ERT2^-GCaMP5G mice (expressing dsRed, or tdTomato and GCaMP5G, under the NG2 promoter, respectively) (see Supplementary Figure 4A for the in vivo setup). Ani9 or its aCSF vehicle were applied to the barrel cortex before CCAO. CCAO resulted in an almost complete cessation of CBF (Figure 7C-D).

**Figure 7:**
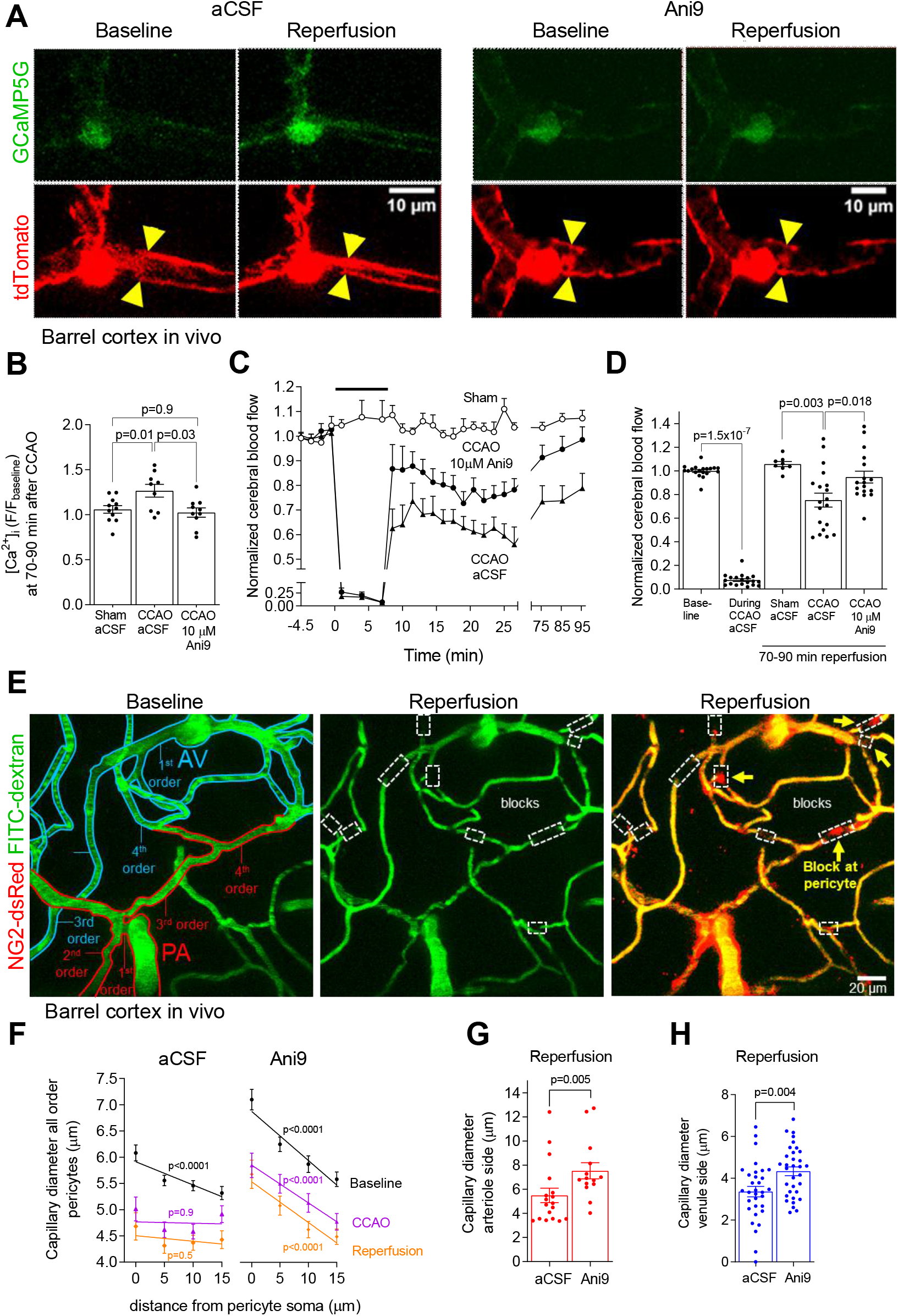
Blocking TMEM16A reduces pericyte contraction and improves CBF post-CCAO. **(A)** In vivo barrel cortex pericytes in P36-P40 NG2-Cre^ERT2^-GCaMP5G mice before and following ~7.5 min CCAO following 1 hr exposure to Ani9 (10 μM) or vehicle. Yellow triangles: sites of changes in vessel diameter. **(B)** Mean GCaMP5G fluorescence (F) in pericyte somata normalized to baseline fluorescence prior to sham-operation or CCAO; each point is mean of data at 70-90 min reperfusion (one-way ANOVA with Tukey’s post hoc test). **(C)** Time course of normalized CBF from laser Doppler flowmetry during sham-operation or CCAO (black bar) in the absence (aCSF) or presence of Ani9 (10 μM) **(D)** Normalized CBF at 70-90 min reperfusion. Points indicate recordings from individual animals (paired two-tailed Wilcoxon test with continuity correction for ‘baseline’ and ‘during CCAO aCSF’ conditions, and one-way ANOVA with Dunnett’s post hoc test for the other conditions) **(E)** In vivo barrel cortex of NG2-dsRed mouse with FITC-dextran in blood. Capillary branching orders are given from the PA or AV. Red and blue tracing indicates vessels on the arteriole or venule side, respectively. White boxes indicate occlusion sites and yellow arrows (right panel) highlight pericytes. **(F)** Mean baseline capillary diameter versus distance from pericyte soma for all branch orders, during CCAO and at 70-90 min reperfusion. Capillary branch orders are 1^st^-3^rd^ order from arteriole: (for aCSF, n=14), (Ani9, n=13); 4^th^-6^th^ order from arteriole or venule: (aCSF, n=4), (Ani9, n=7); or 1^st^-3^rd^ order from venule (aCSF, n=13), (Ani9, n=11). P-value compares the slope of the linear regression line with zero. **(G)** Mean 1^st^ order capillary diameter from PAs at 70-90 min reperfusion in the absence (aCSF) or presence of Ani9 (10 μM). Points indicate individual capillary segments at 0-5 μm from pericyte somata (Mann-Whitney test). **(H)** As for (G) for pericytes on 1^st^-3^rd^ capillary branch order from AVs (unpaired two-tailed Student’s T-test).

Ani9 (10 μM) reduced the CCAO-evoked pericyte [Ca^2+^]_i_ rise after 70-90 min reperfusion (Figure 7A-B), and improved CBF both initially following CCAO (Figure 7C) and after 70-90 min reperfusion (Figure 7C-D). No significant change in CBF was observed in sham-operated mice (Figure 7C), suggesting that the anesthetic did not alter CBF over the duration of the experiments.

Measuring capillary diameter as a function of distance from the pericyte soma at various branch orders from the penetrating arteriole (PA) or ascending venule (AV) (Figure 7E) showed that the CCAO-evoked capillary constriction was largest at the pericyte somata (Figure 7F, aCSF panel). Before CCAO, the capillary diameter was largest at the pericyte somata, as reported above in cortical slices (Figure 3E) and previously in vivo (6, 11), and capillary diameter decreased with distance from the soma, as indicated by the negative slope of the regression line differing significantly from zero (Baseline plots, Figure 7F, aCSF panel). CCAO and reperfusion resulted in a shallower dependence of diameter on distance, with a regression line slope that was no longer significantly different from zero due to constriction of the capillary at the pericyte soma (Figure 7F, aCSF panel). Before CCAO, Ani9 treatment increased the capillary diameter at 0-5 μm from the center of pericyte somata on both the arteriole (p=0.02, 1^st^-3^rd^ order, Mann-Whitney test) and the venule (p=0.0006, 1^st^-3^rd^ order, unpaired two-tailed Student’s T-test) sides of the capillary bed (compare Baseline plots in aCSF and Ani9 panels of Figure 7F), suggesting the existence of some basal TMEM16A activity promoting pericyte tone in cerebral capillaries in vivo (in contrast to the brain slice data above). During reperfusion, Ani9 reduced the capillary constriction on the arteriole (Figure 7G) and the venule (Figure 7H) sides of the capillary bed, and the dependence of diameter on distance from the pericyte soma maintained a steeper negative slope that was significantly different from zero both during and after CCAO (Figure 7F, Ani9 panel). Thus, TMEM16A amplified pericyte-mediated capillary constriction after stroke, and Ani9 prevented this.

### TMEM16A activation enhances ischemia-evoked capillary occlusions at pericyte somata

Stalling of capillary blood flow, in part due to neutrophil block of capillaries, has been reported to decrease CBF both after ischemic stroke and in Alzheimer’s disease (16, 28, 61). We explored whether pericyte-induced narrowing of the capillary lumen during ischemia in vivo could promote transient or permanent stalling of capillary blood flow and whether this was promoted by TMEM16A activity. A FITC-albumin perfusion protocol adapted from David Kleinfeld’s lab (4, 62) was used as a tool to visualize all vessels that had been patent in vivo (see Methods). After sham operation with aCSF (n=4), 7.5 min CCAO with aCSF (n=10) or 7.5 min CCAO with 10 μM Ani9 applied locally to the cortex (n=9), followed by a 1.5 hr period of reperfusion after the CCAO, we introduced FITC-albumin in gelatin into the blood, fixed the brain and imaged sagittal sections of it. CCAO induced capillary blocks and thus evoked a perfusion deficit in the cortex of aCSF-treated mice (Figure 8A; Supplementary Figure 5A-B).

**Figure 8:**
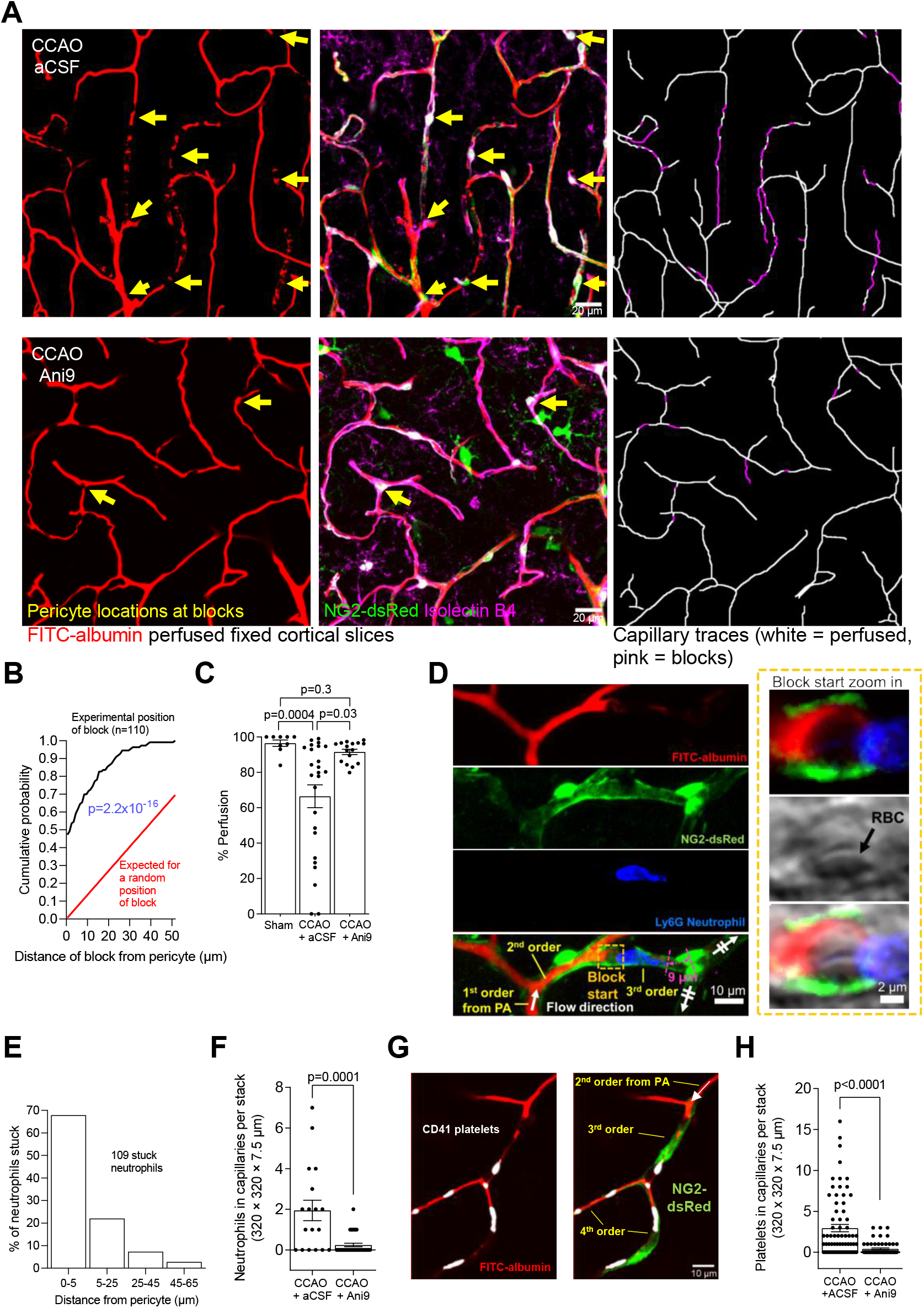
Blocking TMEM16A partially restores capillary perfusion after CCAO. **(A)** Isolectin B4-labelled cortical capillaries in fixed slices (purple); FITC-albumin in gelatin (recolored red) shows perfused vessels 1.5 hrs after sham-operation or CCAO without or with Ani9 (10 μM). Yellow arrows: pericyte somata <5 μm from capillary blocks. Right images show 3D-tracing of FITC-albumin in perfused (white) or unperfused (magenta) capillaries. **(B)** Cumulative probability distribution of distance of 110 pericyte somata to capillary occlusions (black). Red line: predicted distribution assuming pericytes are uniformly spaced along the capillary (see Supplementary Figure 5E) and blocks occur randomly (Kruskal Wallis test with Dunn’s post hoc test). **(C)** Extent of perfusion of 3D traced capillaries. Points indicate individual confocal stacks from P30-72 CCAO (aCSF), P31-83 CCAO Ani9 and P39-62 sham aCSF mice (Kolmogorov-Smirnov test). **(D)** Ly6G-labelled neutrophil in a 3^rd^ order capillary in a fixed cortical slice at 1.5 hours after CCAO. The neutrophil obstructs blood flow (revealed by FITC-gelatin staining). Zoom-in box of right end of the block and left end of neutrophil (“Block start”) shows a possible red blood cell (RBC) to the left of the neutrophil. **(E)** Distribution of 109 neutrophils versus distance from the nearest pericyte soma. **(F)** Neutrophils in cerebral capillaries per confocal stack at 1.5 hours after CCAO in the absence (n=18) or presence (n=33) of Ani9 (10 μM) (Mann-Whitney test). **(G)** CD41-labeled platelets (or aggregates thereof) in 3^rd^ and 4^th^ order capillary branches in a fixed cortical slice at 1.5 hours after CCAO. **(H)** Platelets (or platelet aggregates) in cerebral capillaries per confocal stack at 1.5 hours after CCAO in the absence (n=71) or presence (n=60) of Ani9 (10 μM) (Mann-Whitney test). Number of animals are in Suppl. Table 2.

The cumulative probability distribution for the distance from 110 block sites in capillaries to the nearest pericyte soma, at 1.5 hrs reperfusion, is shown in Figure 8B. The mean distance was 7.2 μm for mice not treated with Ani9. There was 1 pericyte per 148 μm of vessel length (706 pericytes in 104,597 μm of vessel length traced in 10 P30-83 mice: see Supplementary Figure 5E), so if pericytes were uniformly spaced along capillaries and the probability of an occlusion occurring was independent of position in the capillary, then the probability would be uniform from the pericyte soma to half the distance between the pericytes (74 μm, after which the occlusion would be closer to the next pericyte along the capillary). Consequently, the cumulative probability distribution would be a straight line reaching unity at 74 μm from the soma, as shown in Figure 8B. This theoretical distribution differs significantly from the experimentally observed one (p=2.2×10^-16^ from a Kolmogorov-Smirnov test). Thus, the distances between occlusions and pericyte somata are significantly shorter than those expected from a random block of capillaries along their length. This is presumably because the blocks occur close to pericyte somata where capillary constriction is greatest (Figure 7F) owing to most circumferential contractile processes being located near the soma (11). TMEM16A activation, which enhances pericyte contraction after stroke (Figure 7A, EH), will thus increase the occurrence of capillary blocks.

Three-dimensional capillary tracing was used to assess capillary perfusion, as quantified by measuring the length of vessels perfused with FITC-albumin, expressed as a percentage of the total length of isolectin B4 labeled vessels present (Figure 8A). A small perfusion deficit of 3.5% was detected in sham-operated mice, possibly reflecting naturally regressing vessels (63) or artefacts introduced by the perfusion protocol. CCAO increased this deficit to 31% (Figure 8C). Local application of Ani9 reduced the occurrence of blocks (Figure 8A; Supplementary Figure 5A), and reduced the perfusion deficit to 5% (Figure 8C).

To test whether capillary blocks contained neutrophils, we used the neutrophil-specific marker Ly6G, a glycosylphosphatidylinositol–anchored protein, to stain fixed FITC-albumin perfused sagittal sections of mice that underwent CCAO and 1.5 hours of reperfusion after aCSF or Ani9 (10 μM) were applied to the cortical surface (Figure 8D; Supplementary Figure 5B). This revealed a neutrophil density of 1 per 1055 μm of 3D-traced vessel length and a mean neutrophil-to-pericyte distance of 6.8 μm in the cortex of aCSF-treated mice (Supplementary Figure 5E), which is similar to the mean distance of occlusion sites from pericyte somata (see above). Some stalled neutrophils had red blood cells associated with them (Figure 8D). Plotting the distribution of stalled neutrophils as a function of distance from the nearest pericyte showed that 68% of neutrophils were within 5 μm of pericyte somata (Figure 8E). This is consistent with neutrophils becoming trapped in capillaries at contracted pericytes (16, 28, 64) (Figure 8A). Tracking the capillary branching order of neutrophils from the PA or AV in aCSF treated mice, after 1.5 hrs reperfusion, showed that 16% of stalled neutrophils were on 1^st^-2^nd^ order capillary branches from PAs and 41% on 1^st^-2^nd^ order branches from AVs (Supplementary Figure. 5B-D), suggesting that the majority of stalled neutrophils are located in the middle or on the venule side of the capillary bed.

Ani9 treatment significantly lowered the number of neutrophils stalled in capillaries (Figure 8F), reducing the (presumably trapped) neutrophil density 7-fold (to 1 per 7397 μm vessel length; Supplementary Figure 5E). No neutrophils were detected in arterioles, venules or in extravascular areas of the brain parenchyma. Labelling platelets for CD41 revealed that they also tend to occur near sites of stalled capillary blood flow (Figure 8G). Ani9 similarly greatly reduced the number of platelets stalled in capillaries (Figure 8H).

Thus, TMEM16A currents contribute to the ischemia-evoked depolarization of pericytes that facilitates capillary constriction and hence neutrophil and platelet stalling in the acute phase of simulated stroke.

### TMEM16A block improves CBF, reducing infarction and cerebral hypoxia after ischemic stroke in aged mice

To assess whether TMEM16A inhibition confers neuroprotection in a mouse model of stroke which incorporates a major risk factor for stroke, namely aging, we performed ~15 min bilateral CCAO in 15 month old mice followed by 6 hrs reperfusion (see Supplementary Figure 4B for in vivo setup). Similar to the experiments carried out in younger mice, TMEM16A inhibition with Ani9 applied topically to the barrel cortex improved CBF after CCAO as compared to mice treated with aCSF alone (Figure 9A-B). Consistent with this, TMEM16A inhibition reduced infarct size assessed using 2,3,5-triphenyltetrazolium chloride (TTC), which stains (dark red) metabolically active tissue when it is reduced by succinate dehydrogenase (Figure 9C-D). Using pimonidazole (Hypoxyprobe) to measure hypoxia after CCAO in vivo, we found that TMEM16A inhibition significantly decreased hypoxia labeling in the cortex and striatum (Figure 9E-F). Counting the number of hypoxic neurons and glia revealed that the majority (81.2%) of hypoxic cells in the cortex were neurons (Figure 9G-H). Furthermore, loss of cortical NeuN labelling, which is a marker of neuronal injury after ischemic challenge (65), was reduced by TMEM16A inhibition in the cortex of mice undergoing 6 hrs reperfusion (Figure 9I).

**Figure 9:**
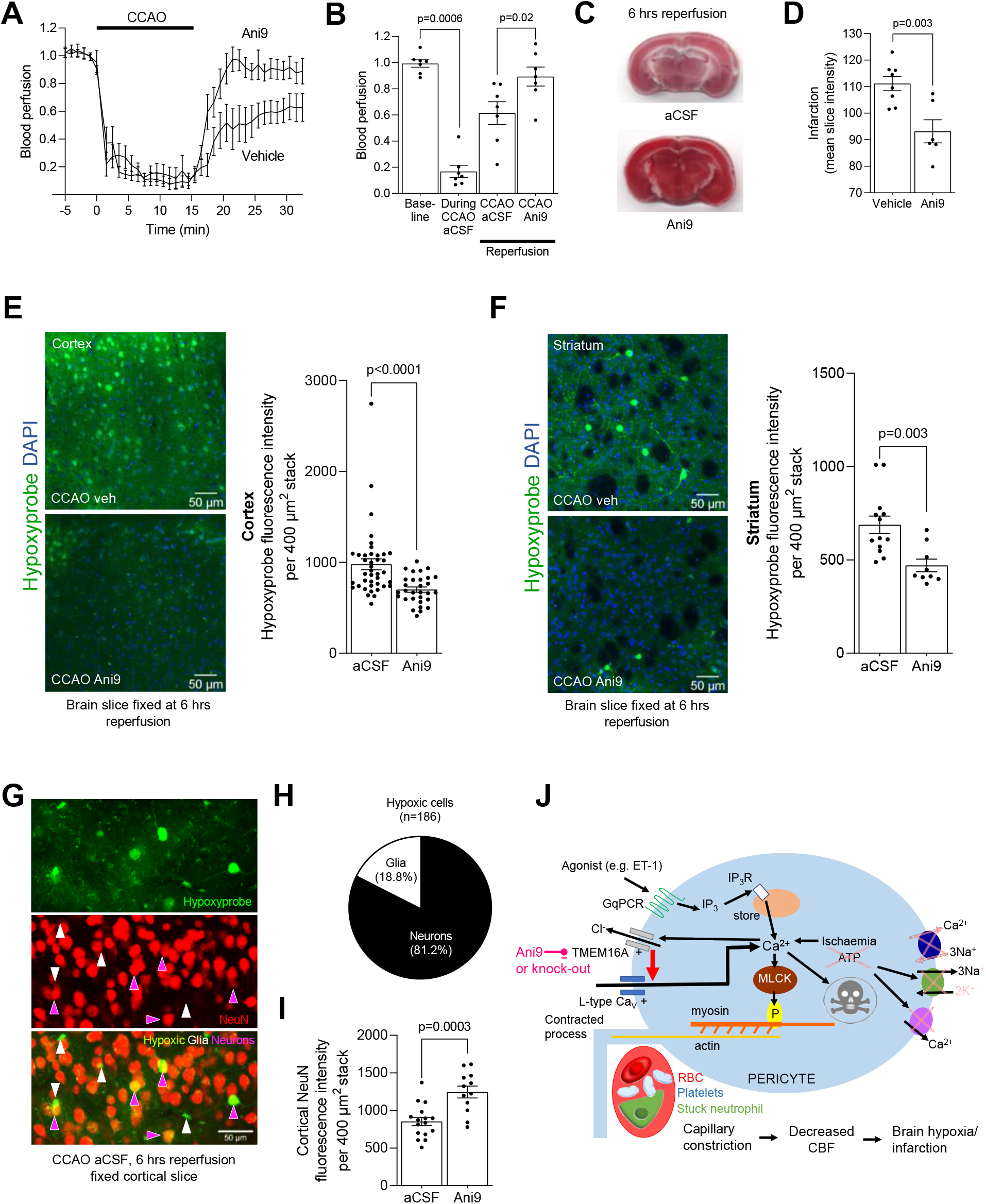
Blocking TMEM16A improves CBF and reduces neuronal hypoxia and infarct size in aged mice. **(A)** Time course of normalized CBF from laser Doppler flowmetry during CCAO (black bar) in the absence (aCSF) or presence of Ani9 (10 μM). **(B)** Normalised CBF during CCAO or reperfusion (average of last 5 minutes of traces in (A)). Points indicate individual CBF recordings from 15 months-old aged mice (paired two-tailed Wilcoxon test with continuity correction and Mann-Whitney test). **(C)** TTC-stained brain sections at 6 hours reperfusion after CCAO in aged mice. **(D)** Infarction quantified from the mean intensity of TTC-stained sections (Mann-Whitney test). Confocal images of fixed cortical **(E)** and striatal **(F)** slices from aged mice that were injected with pimonidazole (Hypoxyprobe) in vivo at 70 minutes after CCAO. Mice underwent 6 hours reperfusion. Bar graphs show Hypoxyprobe intensities in the cortex (E) and striatum (F) from mice treated with aCSF or Ani9 (Mann-Whitney test and unpaired two-tailed Student’s T-test). **(G)** Confocal image of layers II and III in a fixed cortical slice from a mouse undergoing 6 hrs reperfusion after CCAO. **(H)** Proportion of cells with a glial or neuronal morphology labeled with Hypoxyprobe in the cortex quantified from images such as in (H). **(I)** Cortical NeuN fluorescence intensity at 6 hrs reperfusion (Unpaired two-tailed Student’s T-test). **(J)** Schematic of mechanisms revealed. GqPCR activation triggers the IP_3_ pathway; the resulting [Ca^2+^]_i_ rise stimulates TMEM16A and cell depolarisation which promotes Cav-mediated Ca^2+^-entry. In ischemia, low ATP slows Ca^2+^ pumping, leading to TMEM16A activation and initiation of the positive feedback loop that increases [Ca^2+^]_i_, leading to pericyte contraction and death. Neutrophils and platelets become trapped as pericytes contract and capillaries narrow, further lowering cerebral blood flow. TMEM16A inhibition enhances capillary reflow and reduces tissue damage. Animal numbers are in Suppl. Table 2.

Arterial blood pressure rises by ~30% during bilateral common carotid artery occlusion (CCAO) (66, 67), and then slowly recovers to baseline (67). In rodents and humans, low blood pressure has been reported to worsen neurological outcome (presumably by reducing cerebral perfusion: 68, 69). The reduction of brain injury that we observed with TMEM16A inhibition (as assessed by infarct size, hypoxia and loss of neurons: Figure 9) is unlikely to reflect a fall of systemic blood pressure because the TMEM16A inhibitor was applied topically to only the brain parenchyma, where it is expected to decrease local vascular resistance and increase local blood flow, without having a large effect on total vascular resistance and hence on blood pressure. Thus, these data suggest that TMEM16A inhibition has neuroprotective effects that extend beyond the acute phase of ischemic stroke in aged mice.

## DISCUSSION

The key findings of this study are that: (i) Cl^-^ fluxes mediated by the Ca^2+^-gated channel TMEM16A are a crucial determinant of pericyte tone; (ii) TMEM16A is activated during ischemia and evokes a long-lasting pericyte-mediated capillary constriction that reduces CBF and favours neutrophil and platelet stalling; (iii) genetic analysis suggests that increased TMEM16A expression is associated with poor recovery after ischemic stroke (and the genetic proxy for TMEM16A expression was also associated with other psychiatric, neurological and circulatory traits, consistent with its effects on pericyte function and vascular tone). These findings implicate the TMEM16A channel as a new target for ischemic no-reflow, a severe clinical problem with limited available pharmacological treatments.

Pericyte control of capillary diameter is a key determinant of CBF in health and disease (5, 6, 18) (although CBF is, of course, also controlled by adjustment of the diameter of cerebral arterioles). Contrary to expectation, we have shown that pericyte contraction evoked by agonists that release Ca^2+^ from internal stores is largely not triggered directly by the store-released Ca^2+^. Instead, as schematized in Figure 9J, store-released Ca^2+^ activates plasma membrane TMEM16A Cl^-^ channels, and the resulting depolarization caused by Cl^-^ flowing out of the cell activates Cav channels that raise [Ca^2+^]_i_ much more than occurs as a result of the store-released Ca^2+^ alone (Figure 3B, F). This amplification of the agonist-evoked [Ca^2+^]_i_ rise results in a corresponding increase in the capillary constriction produced (Figure 4).

In Ca^2+^-free solution, ET-1 did not trigger a detectable increase in [Ca^2+^]_i_ in pericytes. This suggests that the ET-1-mediated increases in [Ca^2+^]_i_ are small and localized, but they are sufficient to activate TMEM16A. We speculate that this reflects a spatial proximity between sites of Ca^2+^ release and TMEM16A channels in pericytes (70). Deletion of TMEM16A channels from cells expressing smooth muscle myosin heavy chain has been reported to lower vascular resistance and blood pressure (56). Although that work focused mainly on arteries and arterioles, our data suggest that disruption of pericyte contraction may contribute significantly to the vascular function changes following TMEM16A knock-out, particularly in the brain where the majority of the vascular resistance is located in capillaries (3).

Contraction of pericytes generating capillary constriction plays an important role in the decrease of cerebral blood flow that occurs after stroke (6, 18) and in Alzheimer’s disease (11). These conditions generate an initial rise of [Ca^2+^]_i_, which is caused in stroke by a failure of Ca^2+^ pumping and possibly also a release of ET-1 and thromboxane A2 (20, 43), and caused in Alzheimer’s disease by ET-1 release (11). The discovery of extra signaling steps, i.e. activation of TMEM16A channels and membrane depolarization, between this initial [Ca^2+^]_i_ rise and the final constriction of capillaries that leads to neuronal pathology, suggests new opportunities for therapeutic intervention to try to maintain CBF in these conditions. Our data suggest that either directly inhibiting TMEM16A (Figure 6), or else manipulating the [Cl^-^]_i_ gradient to make it become hyperpolarizing rather than depolarizing in pericytes (Figure 4H), are both strategies worth pursuing in order to reduce the deleterious activation of Cav channels that leads to profound capillary constriction, and an ensuing decrease of microvascular blood flow, occlusion of capillaries by neutrophils and platelets, cerebral infarction and hypoxia (Figures 7, 8 and 9). Prolonged CBF decrease eventually leads to pericyte death (6, 23). Pericyte loss leads to BBB breakdown (24–27, 71) which is reduced by inhibiting TMEM16A after stroke (72).

Therapeutic TMEM16A channel inhibitors are not yet available. Our work suggests that Ani9 could form the basis for the design of drugs suitable for use in humans and with adequate BBB permeability. The recently solved cryo-EM structure of the TMEM16A channel, and determination of small molecule binding sites may help this drug discovery effort (73–76). Elucidating the mode of action of Ani9 may aid the design of specific modulators of TMEM16A channel activity for medical therapies to address a range of neurological conditions in which pericytes restrict cerebral blood flow (77, 78).

## METHODS

Detailed descriptions of (i) ex vivo procedures (cortical slice preparation, imaging of capillary diameter and intracellular pericyte Ca^2+^, assessment of cell death, electrophysiology, immunohistochemistry), (ii) in vivo procedures (CCAO, two-photon imaging, cardiac perfusion for 3D capillary tracing), (iii) genetic analyses and (iv) statistics are reported in the Supplemental Material.

### Use of animals

All animal care and experimental protocols were in accordance with the UK Home Office regulations (Guidance on the Operation of Animals, Scientific Procedures Act, 1986 and subsequent modifications). Animal studies are reported in compliance with the ARRIVE guidelines as detailed in the Supplemental Material.

### Human tissue

Human cortical tissue was obtained from patients undergoing brain resection for glioblastoma or thymus cancer as detailed in the Supplemental Material. All work was performed with the informed consent of the patients and ethical approval from the National Health Service (REC number 15/NW/0568 and IRAS project ID 180727).

### Blinding

The experimenters were blind to the condition either during the execution of the experiments and/or during analysis. The methods of analysis were established during study design, and prior to execution of the experiments, to remove possible operator bias.

### Statistics

Statistical tests used are detailed in the Supplemental Material. Bars show the mean ± standard error mean of the individual recordings. P-values <0.05 were considered significant and all tests were 2-tailed.

### Study approval

Animal breeding, experimental procedure and methods of killing were conducted in accordance with the UK Home Office regulations (Guidance on the Operation of Animals, Scientific Procedures Act, 1986). The use of human cortical tissue samples was undertaken under ethical approval from the National Health Service (REC number 15/NW/0568 and IRAS project ID 180727), as detailed in the Supplemental Material.

## AUTHOR CONTRIBUTIONS

N.K., Z.I., D.A., P.T. designed the research; N.K., Z.I., C.P., T.P., P.S., D.G., D.A., P.T. performed experiments and/or analysed data; J.R. generated the floxed TMEM16A mice; H.S. collected and provided human cortical tissue samples; D.A. and P.T. obtained funding and supervised the research; N.K., Z.I., D.G., D.A., P.T. wrote the paper; N.K. and Z.I. are co-first authors; authorship order reflects the fact that N.K. had a uniquely important role in driving key developments in the work. The authors have declared that no conflict of interest exists. D.G. is employed part-time by Novo Nordisk outside of and unrelated to the work in this paper.

## ACKNOWLEDGEMENTS

We thank Frank Kirchhoff (Homburg) for providing the NG2-Cre mouse, Walter Marcotti (Sheffield) for providing the floxed TMEM16A mouse, and Stuart Martin (UCL) for his assistance with the genotyping. Supported by: a BBSRC LIDo PhD studentship to NK, a British Heart Foundation (BHF) postdoctoral post to ZI, a Wellcome Trust Oxion PhD studentship to C.P., an EMBO Fellowship to TP, the BHF Centre of Excellence (RE/18/4/34215) for DG, an ERC Advanced Investigator Award (BrainEnergy) and Wellcome Trust Senior Investigator Award (099222/Z/12/Z) to DA, a BHF project grant (PG/19/8/34168) to PT and Olster Memorial Fund and Physiological Society sabbatical Travel Grants to PT. For the purpose of Open Access, the author has applied a CC BY public copyright licence to any Author Accepted Manuscript version arising from this submission. Graphical abstract was created with BioRender.

## Notes

### Competing Interest Statement

The authors have declared no competing interest.

## REFERENCES

1. Attwell D, et al. Glial and neuronal control of brain blood flow. Nature. 2010;468(7321):232–243.

2. Howarth C, Mishra A, Hall CN. More than just summed neuronal activity: how multiple cell types shape the BOLD response. Philosophical Transactions of the Royal Society of London. 2021;376(1815):20190630.

3. Gould IG, et al. The capillary bed offers the largest hemodynamic resistance to the cortical blood supply. J Cereb Blood Flow Metab. 2017;37(1):52–68.

4. Blinder P, et al. The cortical angiome: an interconnected vascular network with noncolumnar patterns of blood flow. Nat Neurosci. 2013;16(7):889–897.

5. Peppiatt CM, et al. Bidirectional control of CNS capillary diameter by pericytes. Nature. 2006;443(7112):700–704.

6. Hall CN, et al. Capillary pericytes regulate cerebral blood flow in health and disease. Nature. 2014;508(7494):55–60.

7. Rungta RL, et al. Vascular compartmentalization of functional hyperemia from the synapse to the pia. Neuron. 2018;99(2):362–375.

8. Hartmann DA, et al. Brain capillary pericytes exert a substantial but slow influence on blood flow. Nat Neurosci. 2021:24(5):633–645.

9. Gonzales AL, et al. Contractile pericytes determine the direction of blood flow at capillary junctions. Proc Natl Acad Sci U S A. 2020;117(43):27022–27033.

10. Nelson AR, et al. Channelrhodopsin excitation contracts brain pericytes and reduces blood flow in the aging mouse brain in vivo. Front Aging Neurosci. 2020;12:108.

11. Nortley R, et al. Amyloid beta oligomers constrict human capillaries in Alzheimer’s disease via signaling to pericytes. Science (New York, NY. 2019;365(6450):eaav9518.

12. Khennouf L, et al. Active role of capillary pericytes during stimulation-induced activity and spreading depolarization. Brain. 2018;141(7):2032–2046.

13. Zambach SA, et al. Precapillary sphincters and pericytes at first-order capillaries as key regulators for brain capillary perfusion. Proc Natl Acad Sci U S A. 2021;118(26): e2023749118.

14. Ames A, 3rd, et al. Cerebral ischemia. II. The no-reflow phenomenon. Am J Pathol. 1968;52(2):437–453.

15. Hauck EF, et al. Capillary flow and diameter changes during reperfusion after global cerebral ischemia studied by intravital video microscopy. J Cereb Blood Flow Metab. 2004;24(4):383–391.

16. El Amki M, et al. Neutrophils obstructing brain capillaries are a major cause of no-reflow in ischemic stroke. Cell Rep. 2020;33(2):108260.

17. Cho TH, et al. Reperfusion within 6 hours outperforms recanalization in predicting penumbra salvage, lesion growth, final infarct, and clinical outcome. Stroke. 2015;46(6):1582–1589.

18. Yemisci M, et al. Pericyte contraction induced by oxidative-nitrative stress impairs capillary reflow despite successful opening of an occluded cerebral artery. Nat Med. 2009;15(9):1031–1037.

19. Volpe M, Cosentino F. Abnormalities of endothelial function in the pathogenesis of stroke: the importance of endothelin. J Cardiovasc Pharmacol. 2000;35(4 Suppl 2):S45–48.

20. Lampl Y, et al. Endothelin in cerebrospinal fluid and plasma of patients in the early stage of ischemic stroke. Stroke. 1997;28(10):1951–1955.

21. Koudstaal PJ, et al. Increased thromboxane biosynthesis in patients with acute cerebral ischemia. Stroke. 1993;24(2):219–223.

22. Jespersen SN, Ostergaard L. The roles of cerebral blood flow, capillary transit time heterogeneity, and oxygen tension in brain oxygenation and metabolism. J Cereb Blood Flow Metab. 2012;32(2):264–277.

23. Fernandez-Klett F, et al. Early loss of pericytes and perivascular stromal cell-induced scar formation after stroke. J Cereb Blood Flow Metab. 2013;33(3):428–439.

24. Liu Q, et al. Experimental chronic cerebral hypoperfusion results in decreased pericyte coverage and increased blood-brain barrier permeability in the corpus callosum. J Cereb Blood Flow Metab. 2019;39(2):240–250.

25. Armulik A, et al. Pericytes regulate the blood-brain barrier. Nature. 2010;468(7323):557–561.

26. Bell RD, et al. Pericytes control key neurovascular functions and neuronal phenotype in the adult brain and during brain aging. Neuron. 2010;68(3):409–427.

27. Daneman R, et al. Pericytes are required for blood-brain barrier integrity during embryogenesis. Nature. 2010;468(7323):562–566.

28. Erdener SE, et al. Dynamic capillary stalls in reperfused ischemic penumbra contribute to injury: A hyperacute role for neutrophils in persistent traffic jams. J Cereb Blood Flow Metab. 2020; 41:236–252.

29. Hamilton NB, et al. Pericyte-mediated regulation of capillary diameter: a component of neurovascular coupling in health and disease. Front Neuroenergetics. 2010;2:5.

30. Kamouchi M, et al. Calcium influx pathways in rat CNS pericytes. Brain Res Mol Brain Res. 2004;126(2):114–120.

31. Hill-Eubanks DC, et al. Calcium signaling in smooth muscle. Cold Spring Harb Perspect Biol. 2011;3(9):a004549.

32. Caputo A, Caci E, Ferrera L, Pedemonte N, Barsanti C, Sondo E, et al. TMEM16A, a membrane protein associated with calcium-dependent chloride channel activity. Science (New York, NY). 2008;322(5901):590–594.

33. Yang YD, et al. TMEM16A confers receptor-activated calcium-dependent chloride conductance. Nature. 2008;455(7217):1210–1215.

34. Schroeder BC, et al. Expression cloning of TMEM16A as a calcium-activated chloride channel subunit. Cell. 2008;134(6):1019–1029.

35. Bulley S, and Jaggar JH. Cl channels in smooth muscle cells. Pflugers Arch. 2014;466(5):861–872.

36. Hubner CA, et al. Regulation of vascular tone and arterial blood pressure: role of chloride transport in vascular smooth muscle. Pflugers Arch. 2015;467(3):605–614.

37. Pallone TL, Huang JM. Control of descending vasa recta pericyte membrane potential by angiotensin II. Am J Physiol Renal Physiol. 2002;282(6):F1064–1074.

38. Zeisel A, et al. Molecular architecture of the mouse nervous system. Cell. 2018;174(4):999–1014.

39. Kawamura H, et al. Effects of angiotensin II on the pericyte-containing microvasculature of the rat retina. J Physiol. 2004;561(3):671–683.

40. Vanlandewijck M, et al. A molecular atlas of cell types and zonation in the brain vasculature. Nature. 2018;554(7693):475–480.

41. Mishra A, et al. Imaging pericytes and capillary diameter in brain slices and isolated retinae. Nat Protoc. 2014;9(2):323–336.

42. Adomaviciene A, et al. Putative pore-loops of TMEM16/Anoctamin channels affect channel density in cell membranes. J Physiol. 2013;591:3487–3505.

43. Xiao Q, et al. Voltage- and calcium-dependent gating of TMEM16A/Ano1 chloride channels are physically coupled by the first intracellular loop. Proc Natl Acad Sci U S A. 2011;108(21):8891–8896.

44. Faraco G, et al. Circulating endothelin-1 alters critical mechanisms regulating cerebral microcirculation. Hypertension. 2013;62(4):759–766.

45. Oh SJ, et al. MONNA, a potent and selective blocker for transmembrane protein with unknown function 16/anoctamin-1. Molecular Pharmacol. 2013;84(5):726–735.

46. Seo Y, et al. Ani9, A novel potent small-molecule ANO1 inhibitor with negligible effect on ANO2. PLoS One. 2016;11(5):e0155771.

47. Yang AC, et al. A human brain vascular atlas reveals diverse cell mediators of Alzheimer’s disease risk. bioRxiv. 2021:2021.04.26.441262.

48. Ximerakis M, et al. Single-cell transcriptomic profiling of the aging mouse brain. Nat Neurosci. 2019;22(10):1696–1708.

49. Huang W, et al. Novel NG2-CreERT2 knock-in mice demonstrate heterogeneous differentiation potential of NG2 glia during development. Glia. 2014;62:896–913.

50. Mayr D, et al. Characterization of the two inducible Cre recombinase-based mouse models NG2-CreER ^TM^ and PDGFRb-P2A-CreER ^T2^ for pericyte labeling in the retina. Curr Eye Res 2021; doi: 10.1080/02713683.2021.2002910. Online ahead of print.

51. Burgess S, et al. Guidelines for performing Mendelian randomization investigations. Wellcome Open Res. 2019;4:186.

52. Malik R, et al. Multiancestry genome-wide association study of 520,000 subjects identifies 32 loci associated with stroke and stroke subtypes. Nat Genet. 2018;50(4):524–537.

53. Soderholm M, et al. Genome-wide association meta-analysis of functional outcome after ischemic stroke. Neurology. 2019;92(12):e1271–1283.

54. GTEx Consortium, Laboratory Data Analysis Coordinating Center Analysis Working Group, Statistical Methods Analysis Working Group, Enhancing GTEx Groups, NIH Common Fund BCSSN, Biospecimen Collection Source Site RPCI, et al. Genetic effects on gene expression across human tissues. Nature. 2017;550(7675):204–213.

55. Evangelou E, et al. Genetic analysis of over 1 million people identifies 535 new loci associated with blood pressure traits. Nat Genet. 2018:50(10):1412–1425.

56. Heinze C, et al. Disruption of vascular Ca^2+^-activated chloride currents lowers blood pressure. J Clin Invest. 2014;124(2):675–686.

57. Matchkov VV, et al. The role of Ca^2+^ activated Cl^-^ channels in blood pressure control. Curr Opin Pharmacol. 2015;21:127–137.

58. Ettehad D, et al. Blood pressure lowering for prevention of cardiovascular disease and death: a systematic review and meta-analysis. Lancet. 2016;387(10022):957–967.

59. Staley JR, et al. PhenoScanner: a database of human genotype-phenotype associations. Bioinformatics. 2016;32(20):3207–3209.

60. Murphy TH, et al. Two-photon imaging of stroke onset in vivo reveals that NMDA-receptor independent ischemic depolarization is the major cause of rapid reversible damage to dendrites and spines. J Neurosci. 2008;28(7):1756–1772.

61. Cruz Hernandez JC, et al. Neutrophil adhesion in brain capillaries reduces cortical blood flow and impairs memory function in Alzheimer’s disease mouse models. Nat Neurosci. 2019;22(3):413–420.

62. Tsai PS, et al. Correlations of neuronal and microvascular densities in murine cortex revealed by direct counting and colocalization of nuclei and vessels. J Neurosci. 2009;29(46):14553–14570.

63. Schager B, Brown CE. Susceptibility to capillary plugging can predict brain region specific vessel loss with aging. J Cereb Blood Flow Metab. 2020;40(12):2475–2490.

64. Rolfes L, et al. Neutrophil granulocytes promote flow stagnation due to dynamic capillary stalls following experimental stroke. Brain Behav Immun. 2021;93:322–330.

65. Liu F, et al. TTC, fluoro-Jade B and NeuN staining confirm evolving phases of infarction induced by middle cerebral artery occlusion. J Neurosci Methods. 2009;179(1):1–8.

66. Lataro RM, et al. Baroreceptor and chemoreceptor contributions to the hypertensive response to bilateral carotid occlusion in conscious mice. Am J Physiol Heart Circ Physiol. 2010;299:H1990–1995.

67. Kawasaki S et al. Effects of edaravone on nitric oxide, hydroxyl radicals and neuronal nitric oxide synthase during cerebral ischemia and reperfusion in mice. J Stroke Cerebrovasc Dis. 2020;29:104531.

68. Ji X, et al. The effects of blood pressure and urokinase on brain injuries after experimental cerebral infarction in rats. Neurol Res. 2009;31: 204–208.

69. Ouyang M, et al. Low blood pressure and adverse outcomes in acute stroke: HeadPoST study explanations. J Hypertens 2021;39:273–279.

70. Jin X, et al. Activation of Ca^2+^-activated Cl^-^ channel ANO1 by localized Ca^2+^ signals. J Physiol. 2014;594:16–30.

71. Nikolakopoulou AM, et al. Pericyte loss leads to circulatory failure and pleiotrophin depletion causing neuron loss. Nat Neurosci. 2019;22(7):1089–1098.

72. Liu PY, et al. TMEM16A Inhibition Preserves Blood-Brain Barrier Integrity After Ischemic Stroke. Front Cell Neurosci. 2019;13:360.

73. Paulino C, et al. Activation mechanism of the calcium-activated chloride channel TMEM16A revealed by cryo-EM. Nature. 2017;552(7685):421–425.

74. Paulino C, et al. Structural basis for anion conduction in the calcium-activated chloride channel TMEM16A. Elife. 2017;6:e26232.

75. Dang S, et al. Cryo-EM structures of the TMEM16A calcium-activated chloride channel. Nature. 2017;552(7685):426–429.

76. Dinsdale, et. al. An outer-pore gate modulates the pharmacology of the TMEM16A channel. Proc Natl Acad Sci U S A. 2011;108(21):8891–8896.

77. Cheng J, et al. Targeting pericytes for therapeutic approaches to neurological disorders. Acta Neuropathol. 2018;136(4):507–523.

78. Korte N, et al. Cerebral blood flow decrease as an early pathological mechanism in Alzheimer’s disease. Acta Neuropathol. 2020;140(6):793–810.

