## Supplemental information for "The Ca^2+^-gated Cl^-^ channel TMEM16A amplifies capillary pericyte contraction reducing cerebral blood flow after ischemia"

### DETAILED METHODS INFORMATION

#### Animals

Sprague Dawley rats of either sex were group-housed or kept in litters in open-top cages until experimental use for capillary brightfield imaging and electrophysiology at P20-25 (weaning age is P21). Mice were housed in individually ventilated cages in pairs or groups of 3-5. Mice of either sex were used at P32-P116 except for experiments on aging in Figure 9 for which female mice at 15 months of age were used. NG2-dsRed mice expressing a dsRed fluorophore under the NG2 proteoglycan promoter allowed visualisation of cells expressing NG2 including pericytes, smooth muscle cells (SMCs) and oligodendrocyte precursor cells (OPCs). NG2-Cre$^{ERT2}$-GCaMP5G mice were obtained by crossing tamoxifen-inducible NG2-Cre$^{ERT2}$ knock in mice (1) with floxed GCaMP5G-IREStdTomato mice (2). NG2-Cre$^{ERT2}$ TMEM16A (ANO1) knock-out (KO) mice were obtained by crossing NG2-Cre$^{ERT2}$ mice with floxed TMEM16A KO mice (3). These crosses allowed TMEM16A KO or co-expression of the genetically-encoded $Ca^{2+}$ indicator (GCaMP5G) and tdTomato fluorophore (as a morphological marker) in NG2-expressing cells after oral gavage of tamoxifen. Tamoxifen dissolved in corn oil was given at 100 mg/kg body weight by oral gavage once per day for four consecutive days to adult >P21 mice or to lactating mothers once per day at P8-9 (1). All experiments were performed in age and sex matched littermate control mice from 2 weeks after tamoxifen administration. Animals were maintained in a 12 h light/dark cycle, at controlled temperatures (20–23°C) and fed normal chow and untreated tap water ad libitum. Treated Aspen chip and Sizzle-Nest bedding and one tunnel for environmental enrichment were present in each cage. Animals were sacrificed by cervical dislocation for all experiments using acute cortical slices or aged mice (Figures 1-6, 9) or by cardiac perfusion under terminal anesthesia for in vivo experiments (Figures 7-8). Animal breeding, experimental procedure and methods of killing were conducted in accordance with the UK Home Office regulations (Guidance on the Operation of Animals, Scientific Procedures Act, 1986).

**Human tissue**

Human cortical tissue, removed to gain access to underlying tumours in patients, was provided by Huma Sethi, a neurosurgeon at the National Hospital for Neurology and Neurosurgery, Queen Square. The tissue was obtained from five patients: three glioblastoma patients (a 50-year-old male, a 66-year-old male and a 74-year-old female), one thymus cancer patient (a 40-year-old female) and one sarcoma metastasis patient (a 46-year-old female). All work was performed with the informed consent of the patients and ethical approval from the National Health Service (REC number 15/NW/0568 and IRAS project ID 180727).

**Human genetics**

Genetic association summary data for *TMEM16A* expression were obtained from the Genotype Tissue Expression (GTEx) project (gtexportal.org). The version 8 dataset, which included 15,201 samples across 49 tissues from a multi-ethnic group of 838 individuals, was used to identify single-nucleotide polymorphism genetic variants in the *TMEM16A* gene that associated with altered *TMEM16A* expression in any of the tissues considered in the database. Expression data from GTEx version 7, including 10,294 samples across 48 tissues from a multi-ethnic group of 620 individuals, were used to examine the association of variants with pooled *TMEM16A* expression across all tissues. Genome-wide association study (GWAS) summary data for diastolic blood pressure were obtained from a study of 757,601 European ancestry participants (4). GWAS summary data for risk of developing ischemic stroke and recovery after ischemic stroke were obtained from the MEGASTROKE and GISCOME consortia, respectively. The MEGASTROKE study included 60,341 ischaemic stroke cases and 454,450 controls (5). The GISCOME study included 6,021 stroke cases, adjusted for baseline stroke severity (as measured by the National Institutes of Health Stroke Scale), and measured outcome 60-190 days after ischemic stroke using the modified Rankin Scale (a commonly used score for measuring disability after stroke, ranging from no symptoms to death (6)). The GISCOME study analysis consisted of 3,741 patients that had a score of 0-2 (more favourable outcome) and 2,280 patients with a score of 3-6 (less favourable outcome). Full details of all these studies have been published previously (GTEx Consortium, 2017) (4-6). All studies from which data were used for this work had already obtained relevant authorisation, and further ethical approval or participant consent was not required. The identified genetic proxy for TMEM16A activity, the single nucleotide polymorphism rs755016, was further investigated in a phenome-wide association study using the Medical Research Council Integrative Epidemiology Unit Open GWAS Project (7, 8) (https://gwas.mrcieu.ac.uk/, accessed 17 November 2021).

**Cortical slice preparation**

Cortical slices were prepared for immunostaining, patch-clamp electrophysiology, bright-field imaging of capillary diameter, two-photon imaging of capillary diameter and pericyte intracellular calcium concentration ($[Ca^{2+}]_i$), and cell death experiments. Cortical slices (300 μm-thick) were prepared from neurosurgically resected tissue from patients (40-74 year-old), Sprague Dawley rats (P20-25) or NG2-Cre$^{ERT2}$-GCaMP5G mice (P53-116) on a Leica VT1200S vibratome in ice-cold, oxygenated (95% $O_2$/5% $CO_2$) slicing solution (9). The slicing solution contained (in mM) 93 N-methyl-D-glucamine chloride, 2.5 KCl, 30 $NaHCO_3$, 10 $MgCl_2$, 1.2 $NaH_2PO_4$, 25 glucose, 0.5 $CaCl_2$, 20 HEPES, 5 Na-ascorbate, 3 Na pyruvate, 1 kynurenic acid. The slices were incubated at 37°C in the slicing solution (20 min), and then transferred into a modified solution at room temperature in which the NMDG-Cl, $MgCl_2$, $CaCl_2$ and Na ascorbate were replaced by (in mM) 92 NaCl, 1 $MgCl_2$ 2 $CaCl_2$ and 1 Na-ascorbate. Experiments involving cortical slices were performed within 3-4 hrs of sacrificing rodents or arrival of the human tissue from the hospital (transported in oxygenated ice-cold slicing solution within 20 min to the laboratory). For incubations with IB4, which binds α-D-galactose groups in the basement membrane of capillaries (10), human cortical slices were placed for 2 hrs in oxygenated HEPES-buffered solution containing 10 µg/ml IB4 and (in mM): 140 NaCl, 10 HEPES, 2.5 KCl, 1 $NaH_2PO_4$, 10 glucose, 2 $CaCl_2$, 1 $MgCl_2$ (pH=7.4). For immunohistochemistry, human slices were fixed in 4% paraformaldehyde for 30 min at room temperature on a shaker.

**Cortical slice capillary pericyte imaging**

Acute cerebral cortical slices were perfused at a rate of 5 ml/min with heated (~34 °C) artificial cerebrospinal fluid (aCSF) solution containing (in mM) 124 NaCl, 2.5 KCl, 26 $NaHCO_3$, 1 $MgCl_2$, 1 $NaH_2PO_4$, 10 glucose, 1 ascorbate and 2 $CaCl_2$. The aCSF was gassed with 20% $O_2$, 5% $CO_2$, 75% $N_2$. For experiments using TMEM16A KO mice, endothelin-1 (ET-1) was applied following a 5-min baseline. For all other imaging experiments, vasoconstrictors or high $[K^+]_o$ were applied, or ischaemia was induced after 15 mins of incubation with Ani9 (2 µM; Sigma-Aldrich, SML1813), MONNA (5 µM; Sigma-Aldrich, SML09092), nimodipine (3 µM; Sigma-Aldrich, N149), 0 $[Ca^{2+}]_o$ + EGTA (0.1 mM), low $[Cl^-]_o$ (32 mM) and/or bumetanide (40 µM; Cayman, #14630). In the low $[Cl^-]_o$ or 0 $[Ca^{2+}]_o$ solution, the osmolarity was maintained by adding an appropriate amount of sodium methanesulfonate (Na-MES) or $MgCl_2$, respectively. In the high $[K^+]_o$ aCSF solution, [KCl] was elevated to 92.5 mM, and [NaCl] was reduced to 4.25 mM to maintain osmolarity unchanged. The oxygen- and glucose- deprived (OGD) aCSF solution was obtained by replacing glucose with 7 mM sucrose and by equilibrating the solution with 5% $CO_2$ and 95% $N_2$.

Two-photon imaging of pericytes expressing tdTomato and GCaMP5G in NG2-$Cre^{ERT2}$-GCaMP5G mice (P53-P116) was performed using a Zeiss LSM780 microscope with the two-photon laser (Ti:sapphireMai Tai DeepSee, Spectra Physics) tuned to 940 nm. Cortical pericytes on the 1$^{st}$, 2$^{nd}$ and 3$^{rd}$ capillary branch order of penetrating arterioles (PAs) were imaged by acquiring Z-stacks (within 30-100 µm cortical depth, 2 µm step size, 150 to 300 nm pixel size, 1.58 to 6.3 µs pixel dwell time) every minute using a 20 x / 1.0 NA water immersion objective (W Plan-Apochromat, Zeiss). Emitted fluorescence was spectrally divided by a 555-nm dichroic mirror and collected by GaAsP detectors. The power under the objective did not exceed 20 mW. Image processing was performed in FIJI (11). To account for movements in the x- and y-dimensions, image stacks were projected at maximum intensity in the z-dimension and co-registered using the StackReg plugin in FIJI. Changes in GCaMP5G fluorescence were measured by drawing regions of interest (ROIs) over the soma and processes in FIJI and by normalizing each intensity value to the baseline average. The outer diameter of capillaries labelled by tdTomato was obtained by drawing a 7-pixel wide line across the capillary (where circumferential processes spanned the lumen) and by automatically calculating the full width at half maximum of the tdTomato intensity profile by fitting a Gaussian distribution to the fluorescence intensity profile of the line using a custom-written macro script in FIJI.

Patch-clamping (detailed below) and brightfield imaging of capillary pericytes in rat cortical slices was performed at 30-50 µm depth within cortical layers III and IV, and images were recorded using an Olympus BW51 microscope equipped with differential interference contrast (DIC), a 40x water immersion objective, a Coolsnap HQ2 or an IRIS 9TM Scientific CMOS camera CCD camera, and ImagePro Plus (Media Cybernetics) or Metafluor (Molecular Devices) acquisition softwares. Images were acquired every 5 or 30 s, with an exposure time of 50 ms. The pixel size was 160 nm. During imaging, any changes in focus were manually restored by carefully adjusting the focus knob between image acquisitions. Internal capillary diameters were measured by manually placing a measurement line perpendicular to the capillary at pericyte somata (Figure 2A, C) using MetaMorph software.

**Ischaemia pericyte death experiments**

Cortical slices were prepared from Sprague Dawley rats as described above. Slices were transferred into conical flasks under OGD or control (aCSF) conditions in the absence of presence of Ani 9 (2 µM). Solution were gassed with 5% $CO_2$ and 95% $N_2$ (for OGD solutions) or 5% $CO_2$ and 95% $O_2$ (for aCSF solutions) at 37 °C for the entire duration of the experiment. The solutions were supplemented with 7.5 µM of the necrosis marker propidium iodide (PI; Sigma-Aldrich, #81845) and kept in the dark for the entire duration of the experiment. After the 1 hr incubation, the slices were swiftly washed 3 times (using the above incubation solutions in the absence of PI), prior to transfer to 12-well plates for fixation in 4% paraformaldehyde for 1 hr on a rocking shaker. Following 3 washes in PBS, slices were incubated in 10 µg/mL IB4 in blocking buffer (10% (vol/vol) horse serum, 0.3% (vol/vol) Triton X-100, 1.5% (wt/vol) glycine and 1% (wt/vol) bovine serum albumin in PBS) overnight at 4°C on a shaker. Slices were then washed 3 times in PBS before incubation with the nuclear stain DAPI in PBS (1:50,000) on a shaker at room temperature for 1 hr. Following 3 washes in PBS, slices were mounted on microscope slides in ProLong™ Glass Antifade Mountant. Z-stacks (xyz dimensions 292 µm x 292 µm x 10 µm) of capillary pericytes were acquired using Leica TCS SP8 or LSM 700 confocal microscopes with 20x objectives at 1 µm z step size in layer IV starting at a depth of 20-30 µm from the surface of the slice to exclude cells damaged by slicing. Pericytes were identified as being embedded in the IB4-labeled capillary basement membrane with a bump-on-a-log morphology. The fraction of dead pericytes was obtained by dividing the number of PI-containing pericytes by the total number of pericytes (Figure 6E-F) counted using the Cell Counter plugin in FIJI. Imaging was performed in 2-4 slices and 3-4 regions per slice for each animal with the number of image stacks representing the statistical unit.

**Electrophysiology**

Whole-cell patch-clamp recordings were performed on an Olympus BW51 microscope (as for brightfield imaging) using a HEKA EPC9 amplifier, controlled via the built-in analogue-to-digital and digital-to-analogue converter and the Patchmaster version 2x32 software (HEKA Elektronik, Lambrecht, Germany). Pipettes were pulled from borosilicate glass capillary tubes (Harvard Apparatus, UK) using a Narishige PC-10 pipette puller (Narishige, Japan). Pipette tip diameter yielded a resistance of 4-5 MΩ in the working solutions. The series resistance was compensated to achieve a maximal effective series resistance lower than 10 MΩ. Currents were filtered at 2 kHz and sampled at 10 kHz. Liquid junction potentials (LJPs) were calculated (12, 13) and corrected for off-line; these LJPs equalled ~-9 and -13 mV for $Cs^+$ and $K^+$ based solutions (see below), respectively. Families of TMEM16A-CaCC currents (Figure 1 C, D) were elicited in response to 1 s pulses from -100 mV to +100 mV in 10 mV increments, each followed by a 0.5 s step to -60 mV and elicited every 3 s from a holding potential of -40 mV; with all command voltages adjusted for the LJP of -9 mV before display. The whole-cell currents shown in Figure 4E were recorded in response to 2 s voltage ramps from -100 to +100 mV from a holding potential of -40 mV, with all command voltages adjusted for the LJP of -13 mV before display.

Rat cortical slices were prepared as described above and incubated at ~34°C for 20 min in a $Ca^{2+}$-free physiological saline solution containing (in mM): NaCl 145, KCl 5, $MgCl_2$ 1, HEPES 10, glucose 10, pH 7.4) supplemented with collagenase 1A (0.5 mg/ml; Sigma-Aldrich, C5138), protease XIV (0.4 mg/ml; Sigma-Aldrich, P5147) and bovine serum albumin (1.0 mg/ml). Slices were then transferred to the imaging chamber and perfused with heated (~34°C) aCSF solution gassed with 20% $O_2$, 5% $CO_2$, 75% $N_2$. For the recordings of whole-cell TMEM16A currents in pericytes, the intracellular solution contained (in mM): 95 $CsMeSO_4$, 30 CsCl, 10 TEA-Cl, 10 EGTA, 10 HEPES, 0.1 $Na_2GTP$, and 6 $CaCl_2$ (pH 7.2) to obtain 0.25

μM $[Ca^{2+}]_i$. The nominally 0 $Ca^{2+}$ solution was obtained by omitting $CaCl_2$ from the intracellular solution. In some experiments, EGTA was replaced with equimolar HEDTA, and 3.1 mM $CaCl_2$ was used to obtain approximately 1.3 μM $[Ca^{2+}]_i$ (calculated with Patcher's Power tool, Dr Francisco Mendez and Frank Wurriehausen, Max-Planck-Institut für biophysikalische Chemie, Göttingen, Germany). For assessing the physiological magnitude of the TMEM16A current in pericytes, the intracellular solution contained (in mM) 10 NaCl, 100 K-aspartate, 30 KCl, 1 $MgCl_2$, 10 EGTA, 10 HEPES, 0.1 $Na_2GTP$, and 6 or 0 $CaCl_2$ to obtain 0.25 μM and 0 $[Ca^{2+}]_i$, respectively. For action potential recordings in pyramidal cortical neurons, the pipette solutions contained (in mM) 140 K-gluconate, 3 KCl, 2 $MgCl_2$, 2 $Na_2ATP$, 10 HEPES, 0.2 EGTA, pH adjusted to 7.2 with KOH. The osmolarity of each intracellular solution was adjusted to 300 mOsm/Kg with mannitol.

**In vivo ischaemia two-photon imaging, hypoxia measures and TTC staining**

NG2-Cre$^{ERT2}$-GCaMP5G mice (P32-P63), NG2-dsRed mice (P30-P83) or aged wild-type mice (15 months) were anesthetised using urethane (1.55 g/kg given in two doses 15 minutes apart; Sigma-Aldrich, #94300) and anesthesia was confirmed by the lack of a withdrawal reflex to a paw pinch. Body temperature was maintained at 36-37°C using a feedback-controlled heating pad. Eyes were protected from drying by applying polyacrylic acid eye drops (Dr. Winzer Pharma GmbH). The trachea was cannulated and mice were mechanically ventilated with medical air supplemented with oxygen using a MiniVent (Model 845). Sutures were placed around the left and right common carotid arteries. The skull was exposed, slightly thinned using a drill and dried using compressed air. A custom-made headplate was centred over the right barrel cortex, 3 mm laterally from the midline and immediately caudal to the coronal suture. The headplate was attached using superglue gel and mice were head fixed to a custom-built stage. The skull was thinned over the left frontal cortex to attach 1 or 2 laser Doppler flowmetry probes spaced ~ 5mm apart. Cerebral blood flow

(CBF) was measured with an OxyFlo Pro laser Doppler system. Laser Doppler traces were extracted in Matlab 2015b. A craniotomy of approximately 2 mm diameter was performed and the dura was removed. The exposed barrel cortex was superfused for 1 hr with 10 µM Ani9 or aCSF vehicle in HEPES-based aCSF and sealed off with 2% agarose in HEPES-buffered aCSF (containing Ani9 or vehicle) covered by a glass coverslip. All mice in the Ani9 and vehicle groups were age-matched to minimise variability in leptomeningeal anastomoses that occurs with aging (14).

For experiments using 15-month-old mice (Figure 9), bilateral common carotid artery occlusion (CCAO) was performed for 15 mins followed by 6 hrs of reperfusion. Hypoxia was assessed in vivo using pimonidazole HCl Hypoxyprobe (HP2-100, Hypoxyprobe Inc.) kit following the manufacturer's instructions. Pimonidazole (60 mg/kg), which binds to thiols in hypoxic cells, was injected intraperitoneally at 70 minutes of reperfusion. After 6 hours of reperfusion, brains were extracted, cut coronally in half and the frontal portion of the brains were kept in paraformaldehyde for 24 hours prior to sectioning for immunohistochemistry using the FITC-conjugated anti-pimonidazole antibody. Posterior portions of the brains were cut into ~2 mm coronal sections (excluding the cerebellum) and incubated with 1.5% TTC (#T8877, Sigma) in PBS for 30 minutes at 37°C. TTC-stained sections were imaged using a photo scanner and the infarct area was quantified in FIJI.

For in vivo two-photon imaging experiments, while under anesthetic NG2-dsRed mice were injected retro-orbitally with the intraluminal dye fluorescein isothiocyanate–dextran (FITC-dextran, MW 70 kDa, 2.5 mg in 50 µl saline; Sigma-Aldrich, #46945) to measure vessel diameters. Image Z-stacks (2 µm step size, 330-420 nm pixel size, 1.58-2.55 µs pixel dwell time) of vessels were acquired every 1.5 min in cortical layers I-IV using a Newport-Spectraphysics Ti:sapphire MaiTai laser pulsing at 80 MHz, and a Zeiss LSM710 microscope with a 20x water immersion objective (NA 1.0). Fluorescence was evoked using a wavelength of 1000 nm for dsRed and FITC-dextran or 940 nm for GCaMP5G and tdTomato. The mean laser power in the focal plane did not exceed 25 mW. CBF recordings were obtained every 1.5 min between imaging stacks (to avoid interference of the two-photon laser with the laser Doppler signal). Imaging and CBF measures were performed before, during, and for 30 min after ~7.5 min of CCAO and again at 70 min, 80 min and 90 min after CCAO. Sham-operated mice treated with the vehicle underwent the same procedure without CCAO. The CCAO-evoked CBF drop was not significantly different between treatment groups. Image stacks were projected at maximum intensity in the z-dimension and co-registered using the StackReg plugin in FIJI. Internal capillary diameters were measured at 0, 5, 10, 15 and 20 μm from the centre of pericyte somata at defined capillary branch orders from penetrating arterioles or ascending venules by drawing 3-pixel wide lines across the FITC-dextran filled capillary perpendicular to its axis in FIJI. The full width at quarter maximum was automatically computed by fitting a Gaussian distribution to the FITC-dextran fluorescence intensity profile of the line using a custom-written macro script in FIJI. The change in GCaMP5G fluorescence was measured by drawing a region of interest over the soma and by normalizing each intensity value to the baseline average.

**Cardiac perfusion for 3D capillary tracing**

At ~1.5 hrs after CCAO, NG2-Cre$^{ERT2}$-GCaMP5G mice or NG2-dsRed mice from in vivo two-photon imaging were perfused at 9 ml/min with 20 ml of warm (34-37 °C) PBS with heparin (20 IU/ml), followed by 20 ml of warm (34-37°C) 0.25% (w/v) FITC-albumin (Sigma-Aldrich, A9771) in 5% (w/v) gelatin from porcine skin (Sigma-Aldrich, G1890) in PBS (following a protocol similar to refs (15, 16)). After placing mice head down into ice for 30 min, the brains were extracted and drop-fixed in 4% PFA overnight. Following 3 x 5 min washes in PBS, sagittal brain slices 100 μm thick were cut in PBS using a Leica VT1200S vibratome. Slices were used for immunohistochemistry or labelled with 10 μg/ml IB4 in blocking buffer at 4 °C overnight on a shaker (followed by 3 x 5 min washes in PBS and mounting with DAPI) for confocal imaging to assess the length of cortical capillaries perfused with FITC-albumin. For the latter, image stacks (213 µm x 213 µm x 18-38 µm, 2 µm Z step size) from 2-3 slices per mouse were acquired in the cortex using a Zeiss LSM 700 confocal microscope and the lengths of perfused vessel segments were quantified by tracing all FITC-albumin filled vessels in 3D using the Simple Neurite Tracer plugin in FIJI. These traces were overlayed with IB4 and NG2 channels for further tracing to obtain the total vessel length including non-perfused vessel segments that are not filled with FITC-albumin (Figure 8A). The length of FITC-albumin filled vessels was then expressed as a percentage of the total vessel length. The shortest distance of an occlusion site to the centre of a pericyte was measured and the average capillary length between pericytes was obtained by dividing the total vessel length traced by the number of pericytes (counted using the Cell Counter plugin in FIJI). To visualise global brain blood perfusion after CCAO or sham surgery (Supplementary Figure 5A), entire sagittal brain sections were imaged using a Zeiss Axioscan Z1 fluorescence multi-slide scanner with a Plan-Apochromat 10x/0.45 M27 objective. Fluorophore excitation was evoked using a Colibri.2 LED at 355-375 nm, 460-480 nm, 545-565 nm or 615-635 nm wavelengths.

**Immunohistochemistry**

TMEM16A staining was performed on P21 rat and human (40-70 year old) cortical slices. Live human cortical slices were prepared from neurosurgically resected tissue and some tissue was labelled for capillaries using IB4 (ThermoFisher, I32450), as described above. P21 rats were anesthetised with 0.2 mg/g body weight sodium pentobarbital (given intraperitoneally; Dopharma Research B.V.) and transcardially perfused at 9 ml/min with ice-cold 20 ml PBS followed by 20 ml 4% paraformaldehyde. Brains were extracted and fixed in 4% paraformaldehyde overnight at 4°C. Following 3 x 5 min washes in PBS, brain slices 100 µm thick were cut in PBS using a LEICA VT1200S vibratome. Human and rodent slices were treated with 100% methanol for 20 min at -20 °C, washed 3 x 5 min with PBS and blocked overnight at 4°C on a shaker in blocking buffer containing 10% (vol/vol) horse serum, 0.3% (vol/vol) Triton X-100, 1.5% (wt/vol) glycine and 1% (wt/vol) bovine serum albumin in PBS. Slices were incubated in blocking buffer with or without the rabbit polyclonal TMEM16A antibody (1:200 dilution, Abcam, Ab53212) and (for rodent slices only) the mouse monoclonal NG2 antibody (1:500, Abcam, Ab50009) or rabbit polyclonal α-SMA antibody (1:100, Abcam, Ab5694) overnight at 4°C on a shaker. Following 4 x 10 min washes with PBS, slices were incubated in blocking buffer with the secondary antibodies (1:500) anti-rabbit Alexa 488 (A21206) and (for rodent slices only) anti-mouse Alexa 555 (A31570) or anti-rabbit Alexa 555 (A31572). To label capillaries in human slices, incubations with secondary antibodies were supplemented with 10 µg/ml IB4. Slices were washed 4 x 10 min in PBS and mounted with the nuclear stain DAPI. Identical tissue processing was performed with the primary and secondary antibodies omitted to assess autofluorescence in human cortical slices. Image stacks were acquired in the cortex at 1 µm step size using ZEN imaging software and a Zeiss LSM 700 confocal microscope equipped with four diode lasers (405/488/555/639) and 20x and 60x objectives. Colocalization of TMEM16A with NG2 or IB4 used to label pericyte was measured in FIJI. Masks were created from binarised pericyte channels and multiplied by the raw TMEM16A channel. The fluorescence intensity of TMEM16A in pericytes was then expressed as a percentage of the total TMEM16A signal across the field.

Brain sections (100 µm thick) from 15-month old wild-type mice stained for pimonidazole (see above) were mounted and imaged by confocal microscopy to assess the mean fluorescence intensity of the hypoxia labelling or further stained using a guinea pig polyclonal anti-NeuN antibody (1:300, #266004, Synaptic Systems) overnight at 4°C. Following 4 x 10 min PBS washes, slices were incubated with anti-guinea pig Alexa 633 (1:500, A21105) overnight at 4°C, washed 4 x 10 min in PBS and mounted with DAPI. The mean fluorescence intensity of the cortical NeuN staining was quantified and the number of hypoxic cells with a neuronal morphology (with and without NeuN labelling) or glial morphology per stack were counted.

Neutrophil staining using the Alexa Fluor 647 anti-mouse Ly-6G rat monoclonal antibody (2.5 µg/ml in blocking buffer, BioLegend, #127610) was performed on 100 µm thick brain sections (following an overnight incubation in blocking buffer) from NG2-Cre$^{ERT2}$-GCaMP5G mice or NG2-dsRed mice that underwent cardiac FITC-albumin perfusion at ~1.5 hrs after common carotid artery occlusion. Following 4 x 10 min washes in PBS, brain slices were mounted with DAPI for confocal imaging. The shortest distance from the periphery of an intravascular neutrophil to the centre of a pericyte was measured using 3D capillary tracing with the Simple Neurite Tracer plugin in FIJI.

**Reagent and drug preparation**

Stock concentrations of drugs dissolved in DMSO were 50 mM for Ani9 and MONNA and 100 mM for bumetanide. ET-1 (Sigma-Aldrich, E7764) stock was 100 µM in $H_2O$ and U46619 (Cayman, #16450) stock was 28.5 mM in methyl acetate. The final concentration of DMSO or methyl acetate in aCSF solutions never exceeded 0.1% v/v or 0.0007% v/v, respectively. Ani9 required brief warming under continuous stirring to be dissolved in aCSF. The photosensitive L-type voltage-gated $Ca^{2+}$ channel blocker nimodipine was prepared (protected from light) fresh from powder in polyethylene glycol 400 at 4.78 mM (PEG-400; Sigma-Aldrich, #202398). Nimodipine was then further diluted in 15% (2-hydroxypropyl)-β-cyclodextrin (Sigma-Aldrich, H107) in PBS. Urethane stock was prepared in saline at 0.18 g/ml. Tamoxifen stock was dissolved overnight in corn oil at 10 mg/ml on a shaker at room temperature before oral gavage.

### Genetic association analysis

All supporting data for this work are available within the article text, Table 1 and Supplementary Table 1. Statistical analysis was performed using the programme R (version 3.4.2). The genetic proxy for TMEM16A activity was selected as a single-nucleotide polymorphism in the *TMEM16A* gene that: (i) associated with increased expression of *TMEM16A* in any tissue at genome-wide significance after applying a Bonferroni correction for multiple testing of 48 tissues ($p<10^{-9}$), and (ii) had a secondary association with increased diastolic blood pressure ($P<0.05$). Clumping was performed to a pairwise linkage disequilibrium threshold of $r^2<0.001$ using the TwoSampleMR package of R (8). Elevated diastolic blood pressure was used as a secondary trait by which to select the genetic proxy for *TMEM16A* activity due to the established role of the TMEM16A channel in controlling the tone and diameter of peripheral resistance arteries, which are a major determinant of diastolic blood pressure (17). The association of variants with pooled *TMEM16A* expression across all tissues was estimated using fixed-effects meta-analysis (18). For Mendelian randomization, the association of the variant proxying TMEM16A activity with (i) risk of ischemic stroke and (ii) recovery after ischemic stroke were examined. A statistical significance threshold of $p<0.025$ was used, after applying a Bonferroni correction for testing of two outcomes. For exploration of pleiotropy, the PhenoScanner database of genetic associations was searched to explore for potential pleiotropic effects of the identified genetic proxy for TMEM16A activity that could bias Mendelian randomization analysis (19). Given that PhenoScanner mostly contains summary data from GWAS analyses, a genome-wide significance threshold was used for identifying such pleiotropic associations ($P<5x10^{-8}$).

### Group sizes and exclusion criteria

Group size for each specific experiment were equal by design and informed by power calculation conducted at priori. No exclusion criteria were applied with the exception of: (i) in vivo CCAO procedure: experiments in which the mean CBF during CCAO did not drop by >85% of the baseline average were excluded in accord with Grubbs' method (20) and (ii) ex vivo electrophysiological studies: cells used for patch-clamp experiments had seal resistance ≥ 1 GΩ at the start of the experiment. If resistance was lower than this value when the whole-cell configurations was achieved, the experiment was terminated.

**Randomisation and Blinding**

Mice or rats of appropriate age were selected randomly on the day of the experiment. For ex vivo experiments, pericytes (in capillaries of a given order), that were imaged or patch-clamped were randomly selected in isolated brain slices.

The experimenters were blind to the condition either during the execution of the experiments and/or during analysis. The methods of analysis were established during study design, and prior to execution of the experiments, to remove possible operator bias. Furthermore, objective methods of analysis, such as automated analysis procedures (e.g. measurements of current amplitude or capillary diameter), built into the analysis software, were used for each group when possible. The methods of analysis were established during study design and prior to the execution of the experiments.

**Statistics**

Data analyses were performed with Igor Pro 8 (Wavemetrics, OR, USA) for all electrophysiology data and FIJI (ImageJ 1.53c, NIH) and MetaMorph (Molecular Devices, USA) for all imaging data. Statistical tests were performed in R (versions 3.4.2 or 3.6.3) or Prism 6 and 9 (GraphPad Software Inc., CA, USA). As detailed in Supplementary Table 2, N represents the number of animals and n the number of slices, cells, processes, capillary segments, laser Doppler recordings or confocal stacks (for cell death measurements experiments). Bars show the mean ± standard error mean of the individual recordings listed above. Shapiro-Wilk or D'Agostino-Pearson omnibus tests were used to assesses the normality of data. For non-normally distributed data, a non-parametric statistical analysis was performed using the Mann-Whitney U Test (comparing 2 groups, unpaired), the Kolmogorov-Smirnov test (comparing cumulative distributions, unpaired), the Wilcoxon test with continuity correction (comparing 2 groups, paired) or the Kruskal-Wallis test with Dunn's *post hoc* test (comparing >2 groups). For normally distributed data, a parametric test was used: p-values were determined using a homoscedastic (equal variance) or heteroscedastic (unequal variance, with Welch's correction) two-tailed Student's t-test (comparing 2 groups) or one-way ANOVA with Tukey, Bonferroni or Dunnett *post hoc* tests (comparing >2 groups). These procedures include correction for multiple comparisons within each figure panel. To test whether capillary diameter changes with distance from pericyte soma, we assessed whether the slope of the linear regression fit significantly deviated from zero. P-values <0.05 were considered significant and all tests were 2-tailed.

SUPPLEMENTARY FIGURES

**Supplementary Figure 1. Expression of TMEM16A**

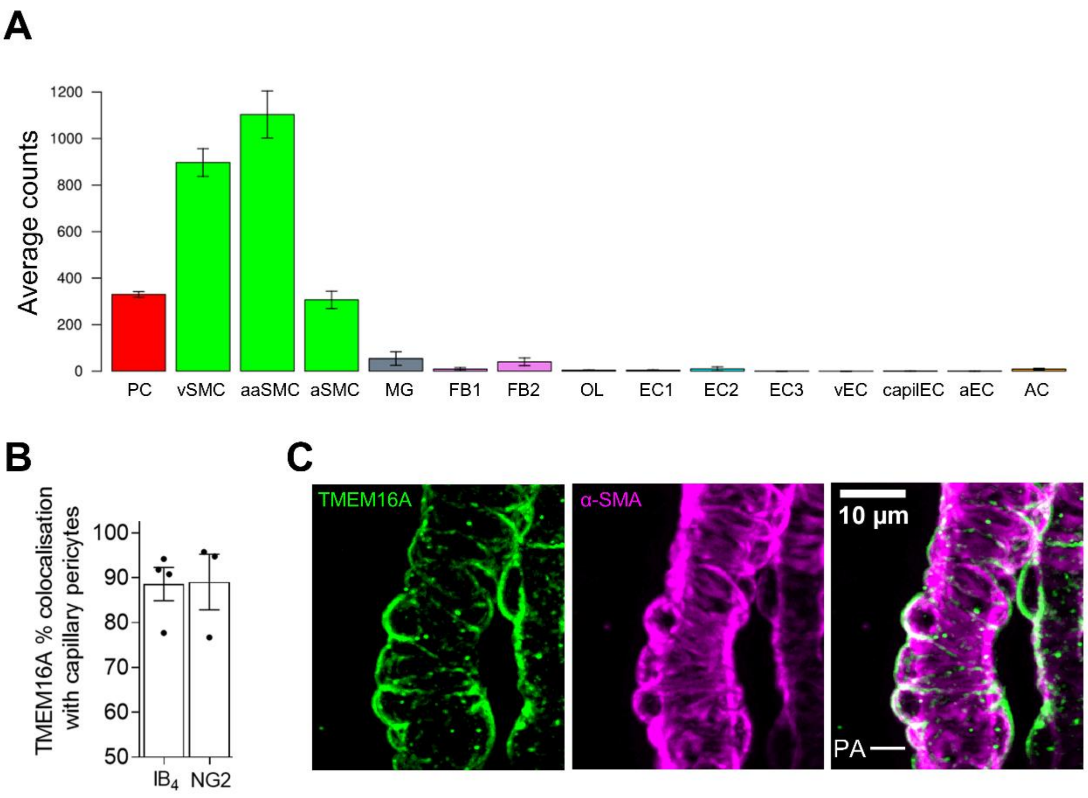


**(A)** Expression of *Tmem16A* (gene name *Ano1*) at the mRNA level in the mouse brain (21) in pericytes (PC), venous smooth muscle cells (vSMCs), arteriolar SMCs (aSMCs), arterial SMCs (aaSMCs), microglia (MG), two fibroblast classes (FB1, FB2), oligodendrocytes (OL), three classes of broadly distributed endothelial cell (EC1-EC3), venous ECs (vEC), capillary ECs (capilEC), arterial ECs, and astrocytes (AC). Data obtained from http://betsholtzlab.org/VascularSingleCells/database.html. **(B)** Percentage of TMEM16A antibody labeling that colocalised with pericytes as defined by their $IB_4$ or NG2 labeling in P21 rat cortical slices (n=3-4). **(C)** Labeling of aSMCs for TMEM16A in P21 rat cortical slices.

**Supplementary Figure 2. Ani9 has no detectable effect on neuronal excitability.**

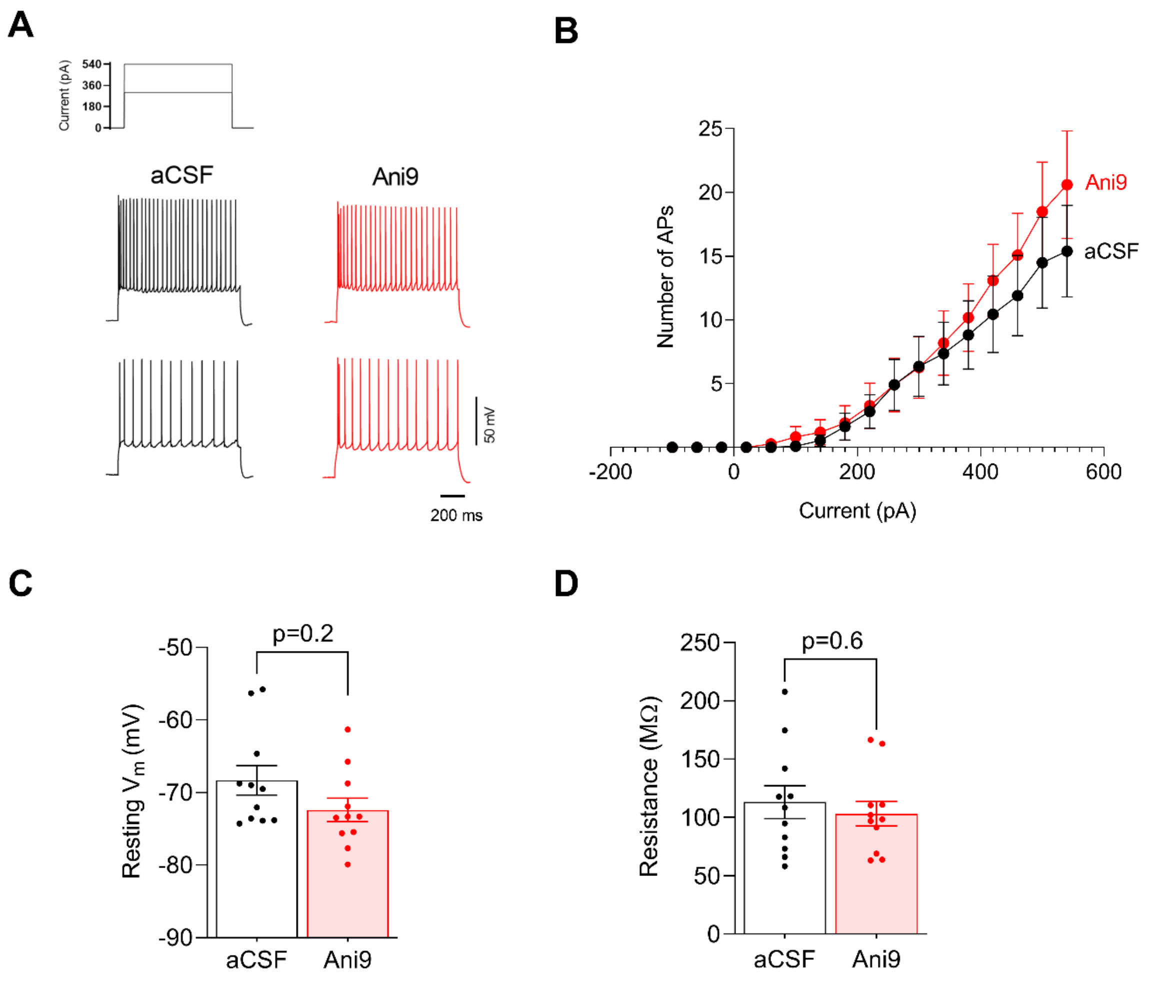


**(A)** Cortical pyramidal neuron action potential response to 1 sec pulses of injected current (inset) in the absence (aCSF) or presence of the TMEM16A blocker Ani9 (2 μM). **(B)** Number of action potentials evoked by the current pulse as a function of the pulse amplitude, measured in the absence (n=10) or presence (n=10) of Ani9 (2 μM) (p=0.3609 at 540 pA of injected current; unpaired Student's t-test). **(C-D)** Resting potential **(C)** and input resistance **(D)** of neurons in the absence (n=10) and presence (n=10) of Ani9 (2 μM) (Mann Whitney test (C), and unpaired two-tailed Student's T-test (D)).

**Supplementary Figure 3. Sex does not affect response to ET-1, OGD, CCAO or Ani9.**

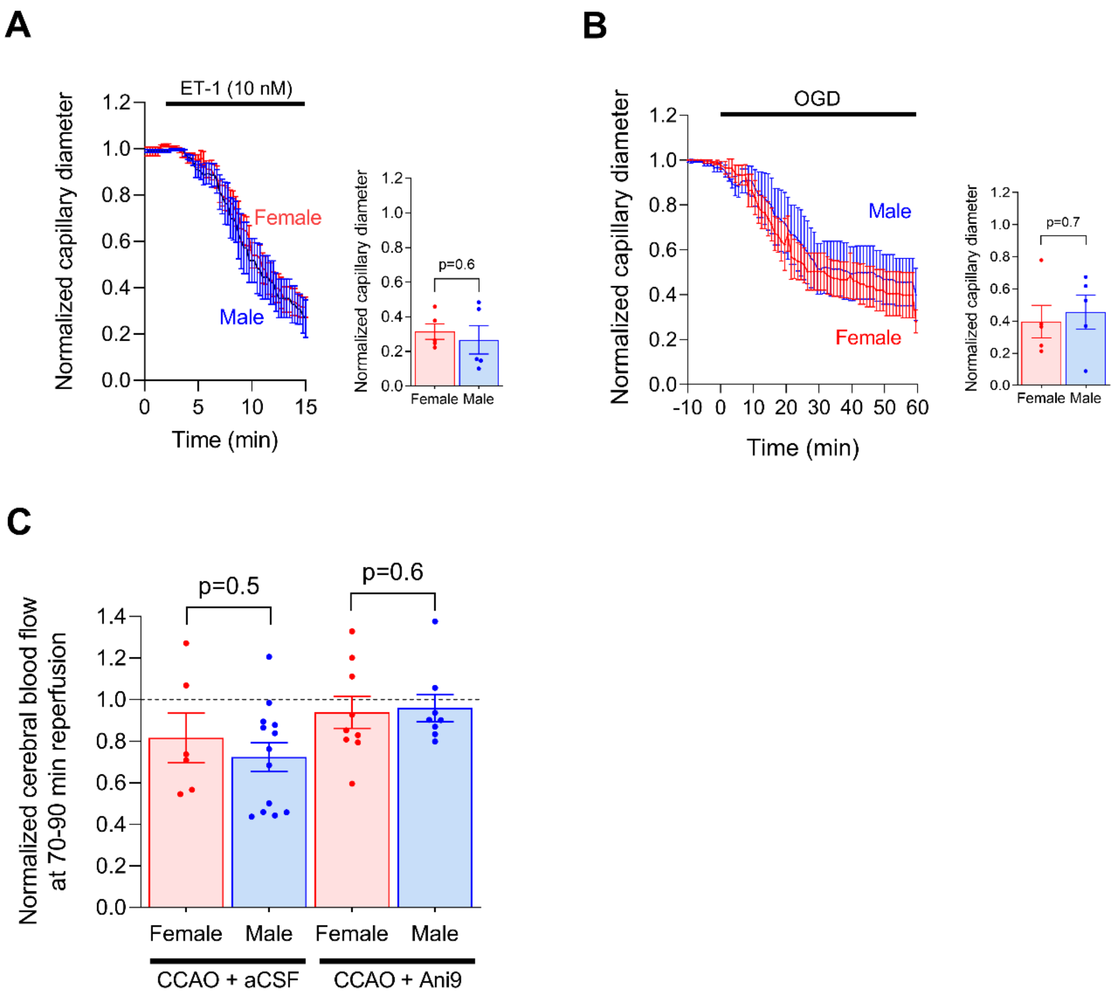


**(A-B)** Response of capillaries in rat brain slices obtained from either male or female rats to ET-1 (A, n=5, male; n=5, female) or OGD (B, n=5, male; n=5 female) (unpaired Student's t-test). Data from both sexes are merged in Figure 2B and Figure 6A **(C)** In vivo response to CCAO without (n=13, male; n=6, female) or with (n=8, male; n=9, female) locally applied Ani9 as in Figure 7C, quantified as normalized CBF after 70-90 mins reperfusion (unpaired, two-tailed Student's t-test and Mann-Whitney test).

**Supplementary Figure 4. Schematic set ups.**

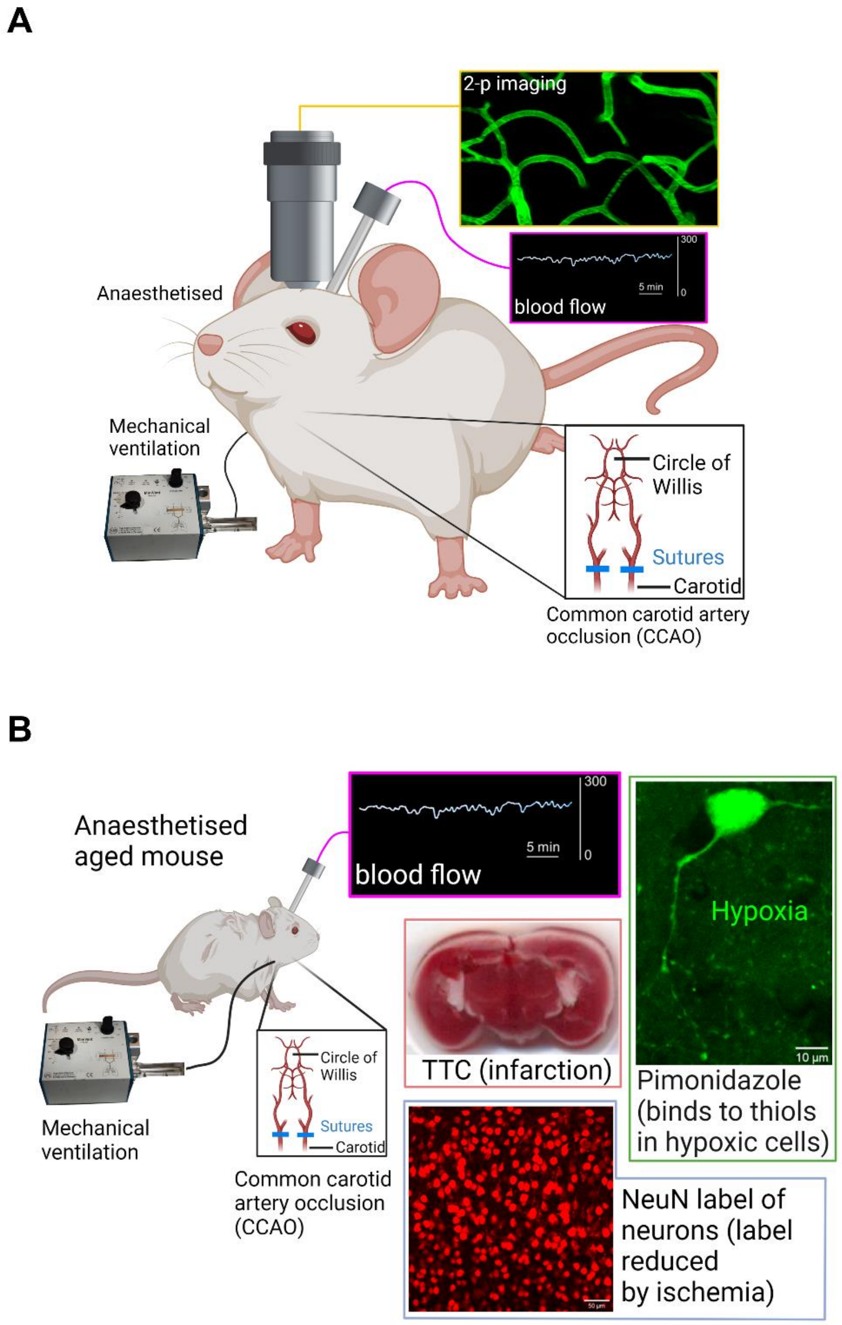


**(A)** Two-photon imaging of capillaries in cortex of anesthetised ventilated mouse, with FITC-dextran in the blood, and laser Doppler measurement of CBF, before and after CCAO to mimic stroke. **(B)** Study of aged mouse using laser Doppler measurement of CBF, and post-CCAO assessment of infarct size (with TTC), hypoxia (with pimonidazole) and loss of NeuN labeling in brain slices taken from the fixed brain. Figures were created with BioRender.com.

**Supplementary Figure 5. Vessel block after CCAO and the neutrophil role in block.**

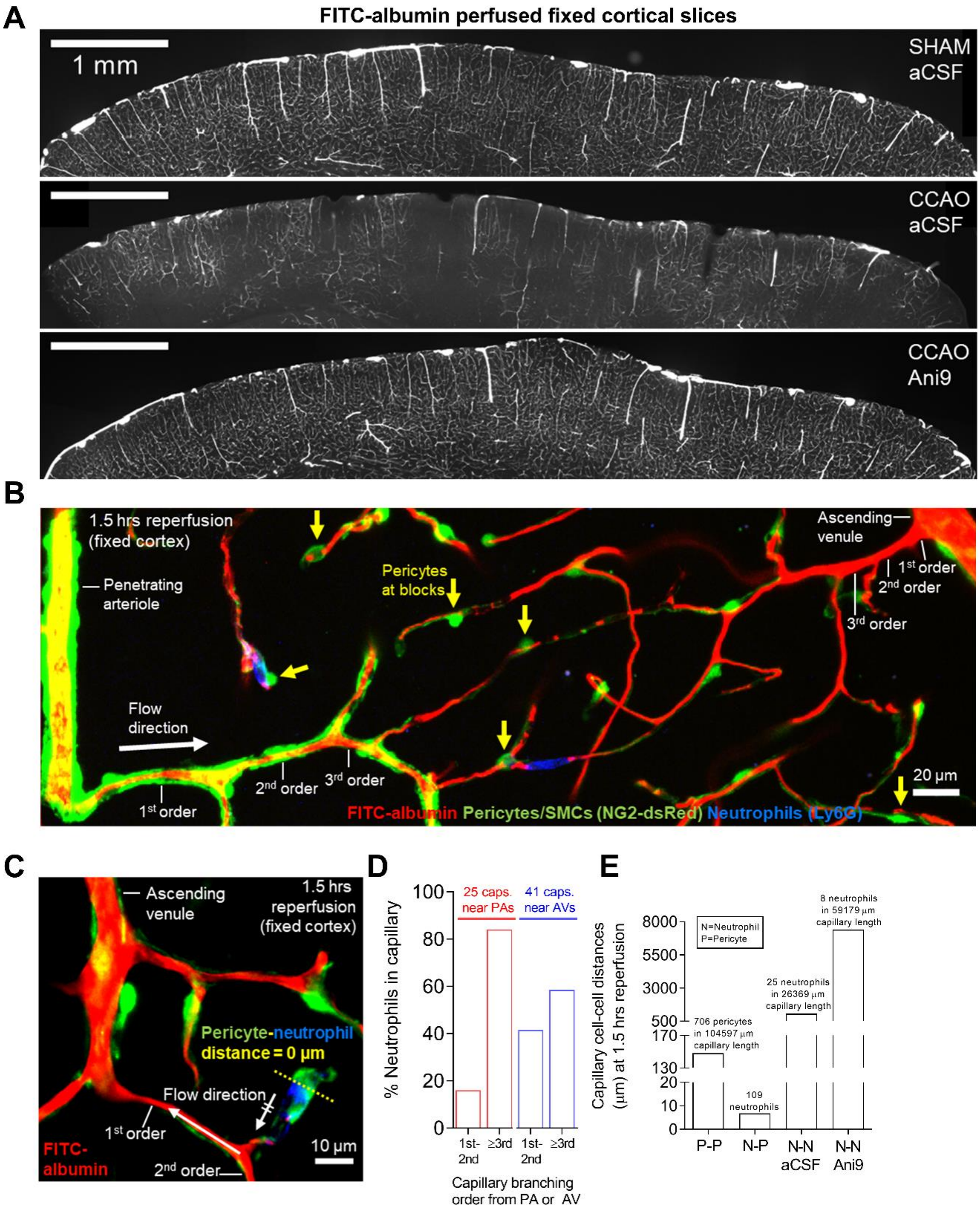


**(A)** Low magnification images of the cortical vasculature with FITC-albumin in the blood, showing that CCAO leads to failure of perfusion of many vessels, and Ani9 ameliorates this. **(B)** Image of an arteriole to venule region with FITC-albumin in the blood (recoloured red), pericytes and SMCs labelled for NG2 (green) and neutrophils labelled with antibody against

Ly6G (blue). CCAO leads to capillary blocks near pericytes which can be associated with neutrophils. **(C)** Example of a stalled capillary near the venule end of the capillary bed with an associated neutrophil. **(D)** Percentage of stalled neutrophils that were in the capillaries of different branch order counting from the penetrating arteriole (PA) or the ascending venule (AV). **(E)** After CCAO and reperfusion, mean distance between pericytes (P-P), from neutrophils stalled in vessels to nearest pericytes (N-P), between neutrophils (N-N), and between neutrophils after Ani9 treatment.

### SUPPLEMENTARY TABLE 1

Supplementary Table 1. Results of the phenome-wide association study investigation of the identified genetic proxy for TMEM16A, rs755016, performed using the Medical Research Council Integrative Epidemiology Unit Open GWAS Project (https://gwas.mrcieu.ac.uk/, accessed 17 November 2021).

| Trait Identifier | Trait | Category | Number of subjects | Effect allele frequency | Beta | Standard error | P-value |
|---|---|---|---|---|---|---|---|
| ubm-a-890 | NODEamps25 0020 | Psychiatric / neurological | 7916 | 0.437 | -0.074 | 0.015 | 1.55E-06 |
| ubm-a-910 | NODEamps100 0019 | Psychiatric / neurological | 7916 | 0.437 | -0.068 | 0.015 | 4.47E-06 |
| ieu-b-39 | Diastolic blood pressure | Circulatory | 703318 | 0.437 | -0.080 | 0.018 | 9.81E-06 |
| ukb-b-7992 | Diastolic blood pressure, automated reading | Circulatory | 436424 | 0.436 | -0.008 | 0.002 | 7.10E-05 |
| ebi-a-GCST007430 | Peak expiratory flow | Lung function | 321047 | 0.437 | -0.010 | 0.003 | 7.67E-05 |
| ukb-b-15085 | Main speciality of consultant (recoded): Adult mental illness | Psychiatric / neurological | 461145 | 0.436 | 0.001 | 0.000 | 9.80E-05 |
| ubm-a-927 | NODEamps100 0036 | Psychiatric / neurological | 7916 | 0.437 | -0.056 | 0.015 | 1.70E-04 |
| ukb-b-3663 | Treatment speciality of consultant (recoded): Adult mental illness | Psychiatric / neurological | 444254 | 0.436 | 0.001 | 0.000 | 2.00E-04 |
| ukb-b-7643 | Diagnoses - secondary ICD10: Z80.3 Family history of malignant neoplasm of breast | Cancer | 463010 | 0.436 | 0.000 | 0.000 | 2.30E-04 |
| ukb-d-H33 | Diagnoses - main ICD10: H33 Retinal detachments and breaks | Opthalmic | 361194 | 0.438 | 0.001 | 0.000 | 2.65E-04 |
| ukb-e-459_AFR | Other disorders of circulatory system | Circulatory | NA | 0.921 | -0.607 | 0.167 | 2.91E-04 |
| ubm-a-915 | NODEamps100 0024 | Psychiatric / neurological | 7916 | 0.437 | -0.054 | 0.015 | 3.02E-04 |
| ukb-d-H7_RETINALDETACH | Retinal detachments and breaks | Ophthalmic | 361194 | 0.438 | 0.001 | 0.000 | 3.05E-04 |
| ukb-e-2020_MID | Loneliness, isolation | Psychiatric / neurological | NA | 0.622 | 0.331 | 0.094 | 4.42E-04 |
| ukb-a-359 | Diastolic blood pressure automated reading | Circulatory | 317756 | 0.437 | -0.009 | 0.003 | 6.23E-04 |
| ebi-a-GCST90000025 | Appendicular lean mass | Anthropometric | 450243 | 0.436 | 0.007 | 0.002 | 6.32E-04 |
| bbj-a-37 | Mean arterial pressure | Circulatory | NA | 0.641 | -0.014 | 0.004 | 6.90E-04 |
| ukb-e-100020_CSA | Typical diet yesterday | Behaviour | NA | 0.664 | 0.466 | 0.137 | 7.00E-04 |
| ubm-a-2313 | NET100 1157 | Psychiatric / neurological | 7916 | 0.437 | -0.054 | 0.016 | 7.08E-04 |
| ubm-a-1348 | NET100 0192 | Psychiatric / neurological | 7916 | 0.437 | -0.054 | 0.016 | 7.59E-04 |
| bbj-a-52 | Systolic blood pressure | Circulatory | NA | 0.641 | -0.013 | 0.004 | 9.81E-04 |

**Supplementary Table 1. Association of traits with the TMEM16A expression proxy rs755016.**

Effect allele is A; non-effect allele is G.

Trait identifier and trait are as defined by https://gwas.mrcieu.ac.uk/

NA in Number of subjects means not available.

Beta: genetic association estimate; SD: standard deviation of beta

P-value: assesses significance of genetic association

### SUPPLEMENTARY TABLE 2 – Number of experiments and animals used in each figure

| Figure | Experiment | # of slices, cells, processes, capillary segments, laser Doppler recordings or confocal stacks (n)/animals (N) |
|---|---|---|
| **1D** | Pericyte patch clamp - mean whole cell TMEM16A current density vs voltage | 1.3 µM $[Ca^{2+}]_i$: n=12, N=8; 0.25 µM $[Ca^{2+}]_i$: n=14, N=9; Ani9: n=9, N=7; 0 $[Ca^{2+}]_i$: n=12, N=7 [n= pericytes, N=rats] |
| **2B** | Effect of Endothelin-1 on capillary diameter | n=10 capillaries, N=8 |
| **2D** | Effect of U46619 on capillary diameter | n=8 capillaries, N=6 |
| **2F** | Endothelin-1 evoked $[Ca^{2+}]_i$ rise in pericytes on $1^{st}$-$3^{rd}$ order capillary branches | $1^{st}$ order: n=13, N=5; $2^{nd}$ order: n=4, N=3; $3^{rd}$ order: n=5, N=3 |
| **3C** | Effect of removing extracellular $Ca^{2+}$ on the Endothelin-1 evoked pericyte $[Ca^{2+}]_i$ rise | Cell n=32, N=3; process n=45, N=3 |
| | | Cell n=32, N=3 |
| | | Process n=79, N=3 |
| **3E** | Effect of removing extracellular $Ca^{2+}$ on the Endothelin-1 evoked capillary constriction | Capillary segments at soma n=40, N=3 |
| | | Capillary segments at 10 µm from soma n=18, N=3 |
| | | Capillary segments at 15-20 µm from soma n=11, N=3 |
| | | Capillary segments at soma n=40; capillary segments 10-20 µm from soma: n=29, N=3 |
| **3G** | Nimodipine effect on Endothelin-1-evoked pericyte $[Ca^{2+}]_i$ rise | Vehicle+ET-1 treated cells ET-1 treated cells n=17, N=3; ET-1 treated cells n=23, N=5; Nimodipine treated cells n=27, N=4 |
| **3G (inset)** | Nimodipine effect on Endothelin-1 evoked pericyte $[Ca^{2+}]_i$ rise in $1^{st}$ vs. $2^{nd}$-$3^{rd}$ order capillary branches | $1^{st}$ order n=9, N=4; $2^{nd}$-$3^{rd}$ order n=10, N=4 |
| **4A** | Capillary diameter with Ani9 and MONNA alone | Ani9: n=13, N=12; MONNA: n=6, N=7 |
| **4B** | Capillary diameter with Ani9 and MONNA + Endothelin-1 | ET-1: n=7, N=6; +Ani9: n=6, N=5; +MONNNA: n=6, N=4 |
| **4C** | Capillary diameter with Ani9 and MONNA + U46619 | U46619: n=8, N=8; +Ani9 n=7, N=7; +MONNNA: n=6, N=5 |
| **4D** | Ani9 effect on ET-1 evoked pericyte $[Ca^{2+}]_i$ rise | ET-1 treated cells n=23, N=5; ET-1 + Ani9 treated cells n=19, N=5 |
| **4E** | Pericyte $I_{MAX}$ | 0 $[Ca^{2+}]_i$ :n=8, N=5; 0.25 $[Ca^{2+}]_i$ :n=10, N=7; 0.25 $[Ca^{2+}]_i$ + Ani9, n=5, N=4 |
| **4F** | Capillary diameter with bumetanide alone | n=9, N=7 |
| **4G** | Capillary diameter with bumetanide and low extracell. $Cl^-$ | Low $Cl^-$: n=7, N=4; Low $Cl^-$ + Bumet: n=8, N=4 |
| **4H** | Capillary diameter with bumetanide and low extracell. $Cl^-$ + Endothelin | ET-1: n=7, N=6; ET-1 + Bumet: n=8, N=3; ET-1 + bumet + Low $Cl^-$: n=7, N=3 |
| **5B** | Effect of TMEM16A KO on Endothelin-1-evoked pericyte contraction | ANO1 KO: n=27, N=5; Wild-type: n=19, N=5 |
| **5D** | Effect of Ani9 on pericyte contraction evoked by high $[K^+]_o$ | With Ani9: n=6, N=5; Without Ani9: n=5, N=5 |
| **6A & B** | Capillary diameter with ACSF, OGD, and OGD + Ani9 | aCSF+Ani9: n=6, N=5; OGD: n=10, N=8; OGD+Ani9: n=6, N=6 |

| | | |
|---|---|---|
| **6D** | Ani9 effect on OGD-evoked pericyte $[Ca^{2+}]_i$ rise | OGD treated cells n = 14, N=3; OGD + Ani9 treated cells n=17, N=3 |
| **6F** | Ani9 effect on OGD-induced cell death (propidium iodide) | aCSF: n=32, N=6; OGD: n=33, N=6; OGD+Ani9: n=35, N=6 [n=image stacks, N=rats] |
| **7B** | Ani9 effect on CCAO-evoked pericyte $[Ca^{2+}]_i$ rise | Sham n=11, N=4; CCAO n=9, N=5; CCAO+Ani9 n=10, N=4 |
| **7C** | Ani9 effect on CBF during CCAO | CCAO n=19, N=10; CCAO+Ani9 n=17, N=9 |
| **7D** | CCAO-evoked drop in CBF | CCAO n=19, N=10 |
| | Ani9 effect on CCAO-evoked drop in CBF at 70-90 min reperfusion | Sham, n=8, N=4; CCAO, n=19, N=10; CCAO+Ani9, n=17, N=9 |
| **7F** | Capillary (all orders) diameter change with distance from pericyte soma | CCAO n=30 pericytes, N=5; CCAO+Ani9 n=32 pericytes, N=5 |
| | Effect of Ani9 on venule side capillary (1st-3rd order) diameter at pericyte somata (0-5 µm) in the normal brain in vivo | Baseline n=32, N=5; Baseline+Ani9 n=33, N=4 |
| | Effect of Ani9 on arteriole side capillary (1st-3rd order) diameter at pericyte somata (0-5 µm) in the normal brain in vivo | Baseline n=36, N=4; Baseline+Ani9 n=35, N=5 |
| **7G** | Effect of Ani9 on arteriole side capillary ($1^{st}$ order) diameter at pericyte somata (0-5 µm) at 70-90 min reperfusion | CCAO, n=18, N=4; CCAO+Ani9 n=14, N=4 |
| **7H** | Effect of Ani9 on venule side capillary ($1^{st}$-$3^{rd}$ order) diameter at pericyte somata (0-5 µm) at 70-90 min reperfusion | CCAO, n=32, N=4; CCAO+Ani9 n=33, N=4 |
| **8B** | Effect of Ani9 on percentage perfusion at 1.5 hrs reperfusion | CCAO, n=25, N=10; CCAO+Ani9 n=16, N=9; Sham, n=9, N=4 |
| **8C** | Cumulative probability of block as a function of distance from pericyte | CCAO, n=110, N=10 |
| **8F** | Effect of Ani9 on neutrophils in capillaries | CCAO, n=18, N=5; CCAO+Ani9, n=33, N=5 |
| **8H** | Effect of Ani9 on platelets in capillaries | CCAO, n=71, N=5; CCAO+Ani9, n=60, N=5 |
| **9A** | Ani9 effect on CBF during CCAO | CCAO n=7, N=4; CCAO+Ani9 n=7, N=4 |
| **9B** | CCAO-evoked drop in CBF | CCAO n=7, N=4 |
| | Ani9 effect on CCAO-evoked drop in CBF at reperfusion | CCAO n=7, N=4; CCAO+Ani9 n=7, N=4 |
| **9D** | Ani9 effect on CCAO-evoked infarction | CCAO n=8, N=4: CCAO+Ani9 n=6, N=3 |
| **9E** | Effect of Ani9 on hypoxia in cortex | CCAO n=40, N=4, CCAO+Ani9 n=30, N=3 |
| **9F** | Effect of Ani9 on hypoxia in striatum | CCAO n=13, N=4, CCAO+Ani9 n=9, N=3 |
| **9I** | Effect of Ani9 on NeuN fluorescence intensity in cortex | CCAO n=16, N=4, CCAO+Ani9 n=12, N=3 |
| **S2B** | Effect of Ani9 on number of action potentials | aCSF: n=10, N=10; Ani9: n=10, N=10 |
| **S2C** | Effect of Ani9 on resting $V_m$ (current clamp) | ACSF: n=11, N=9; Ani9: n=11, N=10 |
| **S2D** | Effect of Ani9 on resistance (current clamp) | ACSF: n=11, N=9; Ani9: n=11, N=10 |
| **S3A** | Effect of ET-1 (10 nM) on capillary diameter | Female: n=5, N=4 ; Male: n=5, N=4 |
| **S3B** | Effect of OGD on capillary diameter | Female: n=5, N=4; Male: n=5, N=4 |

| | | |
|---|---|---|
| **S3C** | Effect of mouse sex on CBF at 70-90 min reperfusion | Female ACSF: n=6, N=3; Male aCSF: n=13, N=7 |
| | | Female Ani9: n=9, N=5; Male Ani9: n=8, N=4 |

### REFERENCES


1. Huang W, et al. Novel NG2-CreERT2 knock-in mice demonstrate heterogeneous differentiation potential of NG2 glia during development. Glia. 2014;62(6):896-913.

2. Gee JM, et al. Imaging activity in neurons and glia with a Polr2a-based and cre-dependent GCaMP5G-IRES-tdTomato reporter mouse. Neuron. 2014;83(5):1058-1072.

3. Amjad A, et al. Conditional knockout of TMEM16A/anoctamin1 abolishes the calcium-activated chloride current in mouse vomeronasal sensory neurons. J Gen Physiol. 2015;145(4):285-301.

4. Evangelou E, et al. Genetic analysis of over 1 million people identifies 535 new loci associated with blood pressure traits. Nat Genet. 2018;50(10):1412-1425.

5. Malik R, et al. Multiancestry genome-wide association study of 520,000 subjects identifies 32 loci associated with stroke and stroke subtypes. Nat Genet. 2018;50(4):524-537.

6. Soderholm M, et al. Genome-wide association meta-analysis of functional outcome after ischemic stroke. Neurology. 2019;92(12):e1271-1283.

7 Elsworth B, et al. The MRC IEU OpenGWAS data infrastructure. bioRxiv 2020;2020.08.10.244293v1. doi: 10.1101/2020.08.10.244293

8. Hemani G, et al. The MR-base platform supports systematic causal inference across the human phenome. Elife (2018) 7:e34408

9. Hall CN, et al. Capillary pericytes regulate cerebral blood flow in health and disease. Nature. 2014;508(7494):55-60.

10. Peters BP, Goldstein IJ. The use of fluorescein-conjugated Bandeiraea simplicifolia B4-isolectin as a histochemical reagent for the detection of alpha-D-galactopyranosyl groups. Their occurrence in basement membranes. Exp Cell Res. 1979;120(2):321-334.

11. Schindelin J, et al. Fiji: an open-source platform for biological-image analysis. Nat Methods. 2012;9(7):676-682.

12. Barry PH, Lynch JW. Liquid junction potentials and small cell effects in patch-clamp analysis. J Memb Biol. 1991;121(2):101-117.

13. Neher E. Correction for liquid junction potentials in patch clamp experiments. Methods in Enzymol. 1992;207:123-131.

14. Hecht N, et al. Cerebral hemodynamic reserve and vascular remodeling in C57/BL6 mice are influenced by age. Stroke. 2012;43(11):3052-3062.

15. Blinder P, et al. The cortical angiome: an interconnected vascular network with noncolumnar patterns of blood flow. Nat Neurosci. 2013;16(7):889-897.

16. Tsai PS, et al. Correlations of neuronal and microvascular densities in murine cortex revealed by direct counting and colocalization of nuclei and vessels. J Neurosci. 2009;29(46):14553-14570.

17 Heinze C, et al. Disruption of vascular $Ca^{2+}$-activated chloride currents lowers blood pressure. J Clin Invest. 2014;124:675-686.

18. GTEx Consortium, Laboratory Data Analysis Coordinating Center Analysis Working Group, Statistical Methods Analysis Working Group, Enhancing GTEx Groups, NIH Common Fund

BCSSN, Biospecimen Collection Source Site RPCI, et al. Genetic effects on gene expression across human tissues. Nature. 2017;550(7675):204-213.

19. Staley JR, et al. PhenoScanner: a database of human genotype-phenotype associations. Bioinformatics. 2016;32(20):3207-3209.

20. Grubbs FE. Procedures for detecting outlying observations in samples. Technometrics. 1969; 11(1):1-21.

21. Vanlandewijck M, et al. A molecular atlas of cell types and zonation in the brain vasculature. Nature 2018;554:475-480.
